# NAD^+^ reduction in glutamatergic neurons triggers fatty acid catabolism and neuroinflammation in the brain, mitigated by SARM1 deletion

**DOI:** 10.1101/2025.05.05.652246

**Authors:** Zhen-Xian Niou, Sen Yang, Andrea Enriquez, Anoosha Sri, Caliel Hines, Jason M. Tennessen, Chia-Shan Wu, Jui-Yen Huang, Hui-Chen Lu

## Abstract

The importance of NAD^+^ homeostasis for neuronal health has been emphasized by studies on nicotinamide mononucleotide adenylyl transferase 2 (NMNAT2), a NAD^+^-synthesizing enzyme, and sterile alpha and TIR motif-containing protein 1 (SARM1), a NAD**^+^** hydrolase. NMNAT2 declines caused by neurodegenerative insults activate SARM1 to degenerate axons. To elucidate the impact of the NMNAT2-SARM1 axis on brain energy metabolism, we employed multi-omics approaches to investigate the metabolic effects caused by neuronal NMNAT2 loss. The loss of NMNAT2 in glutamatergic neurons results in a striking metabolic shift in the cerebral cortex from glucose to lipid catabolism, reduced lipid abundance, and pronounced neurodegenerative phenotypes. Proteomic analysis found that neuronal NMNAT2 loss altered levels of glial enzymes central to glucose and lipid metabolism. Genetic deletion of SARM1 in NMNAT2-deficient mice restores lipid metabolism and mitigates neurodegeneration. Taken together, we show that neuronal NAD^+^ reduction leads to SARM1-dependent maladaptive adaptations in both neurons and glia.

## Introduction

Under normal circumstances, the brain primarily depends on glucose for energy production ^1–3^. As a result, deficits in glycolysis or mitochondrial pyruvate oxidation can severely impair brain function and lead to neurodegenerative conditions ^4–8^. Nicotinamide adenine dinucleotide (NAD^+^), a key cofactor that dictates the rate of glycolytic flux, serves as a vital electron acceptor and is essential for maintaining glucose-derived ATP production. Thus, the enzymatic pathways governing NAD^+^ abundance are crucial in maintaining normal ATP levels to support brain function.

Consistent with the essential role of NAD metabolism in brain health, nicotinamide mononucleotide adenylyl transferase 2 (NMNAT2) and sterile alpha and TIR motif-containing protein 1 (SARM1), two enzymes involved in regulating steady-state NAD^+^ levels, have been recently implicated in a wide range of neuronal disorders ^9–12^. In neurons, NMNAT2 is the most abundant of the three NMNAT isoforms (NMNAT1-3), synthesizing NAD^+^ from nicotinamide mononucleotide (NMN) via the salvage pathway ^12^. We and others have shown that NMNAT2 is critical to maintaining the NAD^+^/NADH redox potential and the survival of long-range cortical axons in mice ^13–15^, while it is dispensable for axonal outgrowth ^13^. Moreover, NMNAT2 overexpression offers neuroprotection in various preclinical models of neurodegeneration and nerve injury ^16–20^. Conversely, SARM1 functions as a NAD hydrolase, and its activation leads to NAD^+^ depletion and axon degeneration^21,22^. SARM1 is a metabolic sensor activated when the ratio of NMN to NAD^+^ increases ^23^. Under normal conditions, SARM1 is kept in an inactive state because NMNAT2 converts NMN to NAD^+^ and maintains low NMN levels ^11,22,24–27^. However, following axonal injury or under pathological conditions, NMNAT2 levels decrease, which subsequently increases NMN and activates SARM1 to trigger axon degeneration processes. Deleting SARM1 protects against the axonopathy phenotypes resulting from NMNAT2 deficiencies ^13,25,28,29^.

While the NMNAT2-SARM1 axis is well-characterized in maintaining NAD redox homeostasis and preserving axonal integrity, its broader impact on brain energy metabolism remains poorly understood. Our recent study demonstrated that neuronal NMNAT2 deficiency leads to a glycolysis deficit in distal axons, accompanied by significant neurodegenerative phenotypes^13^. These findings showed that NMNAT2 loss could disrupt global brain energy metabolism and contribute to neuropathology. Substantial evidence indicates that the CNS exhibits metabolic flexibility to utilize alternative fuels, such as lipids, when glucose availability is limited or glucose metabolism is disrupted ^2,30–38^. We hypothesized that the disrupted NAD^+^ homeostasis caused by neuronal NMNAT2 loss exerts broader impacts by reprogramming brain energy metabolism not only in neurons but also in their neighboring cells, subsequently triggering maladaptive consequences.

In this study, we aimed to define the metabolic consequences of neuronal NMNAT2 loss in the intact brain and determine whether SARM1 deletion could rescue the metabolic aberrations resulting from NMNAT2 deficiency. To this end, we combined metabolomics approaches with proteomics and lipidomic analyses to investigate how neuronal NMNAT2 loss alters energy metabolism. We discovered that the loss of neuronal NMNAT2 leads to a significant reduction in glycolysis and an increase in fatty acid catabolism. Quantitative lipidomic profiling further revealed a substantial depletion of sphingolipids, glycerophospholipids, and fatty acids. This metabolic reprogramming was accompanied by robust neuroinflammation and motor behavior deficits. Complete SARM1 deletion mitigates some but not all of the metabolic alterations caused by neuronal NMNAT2 loss. Our findings reveal that neuronal NMNAT2 plays a crucial role in maintaining NAD homeostasis, and its loss shifts energy metabolism from glucose to lipid usage in the brain.

## Results

### Loss of NMNAT2 from glutamatergic neurons alters cortical energy metabolism

To investigate the impact of NMNAT2 loss on energy homeostasis in the cortex, metabolomic profiling was performed with the P16-P21 cortices of cortical neuron-specific NMNAT2 knockout (cKO) mice and their littermate controls. cKO mice were generated by crossing NMNAT2 conditional allele mice (NMNAT2^f/f^) ^15^ with a Nex-Cre transgenic mouse line, which expresses Cre recombinase in post-mitotic glutamatergic neurons mainly in the cortex and hippocampus from embryonic day 11.5 ^39^. Both gas and liquid chromatography-mass spectrometry (GC-MS and LC-MS) metabolomic analyses were conducted, with LC-MS focusing on redox metabolites and nucleotides because of the known contributions of NAD^+^ to these metabolic pathways ^40^. Partial Least Squares Discriminant Analysis (PLS-DA) of the metabolomic data shows distinct profiles between NMNAT2 cKO (Nex^cre/+^;NMNAT2^f/f^) and control (Nex^cre/+^;NMNAT2^f/+^) cortices, independent of sex (Fig. 1A). Significant decreases in nicotinamide (Fig. 1B) and NAD/NADH ratios in cKO cortices (Fig. 1D) underscore the critical role of NMNAT2 in maintaining cortical NAD^+^ homeostasis. Notably, cKO cortices contain significantly higher levels of sugar derivatives, oxidized glutathione, and purine (adenine and guanine) and pyrimidine (uracil, thymine, and cytosine) nucleotides compared to controls (Fig. 1B). Metabolite Set Enrichment Analysis (MSEA), conducted using MetaboAnalyst 4.0 ^41^, further indicated significant alterations in several pathways: glycolysis, amino acid metabolism, nucleotide sugar metabolism, purine/pyrimidine metabolism, glutathione metabolism, and starch/sugar metabolism (Fig. 1C). These metabolic changes suggest that the loss of neuronal NMNAT2 results in marked impairments in several metabolic pathways.

**Fig. 1.**
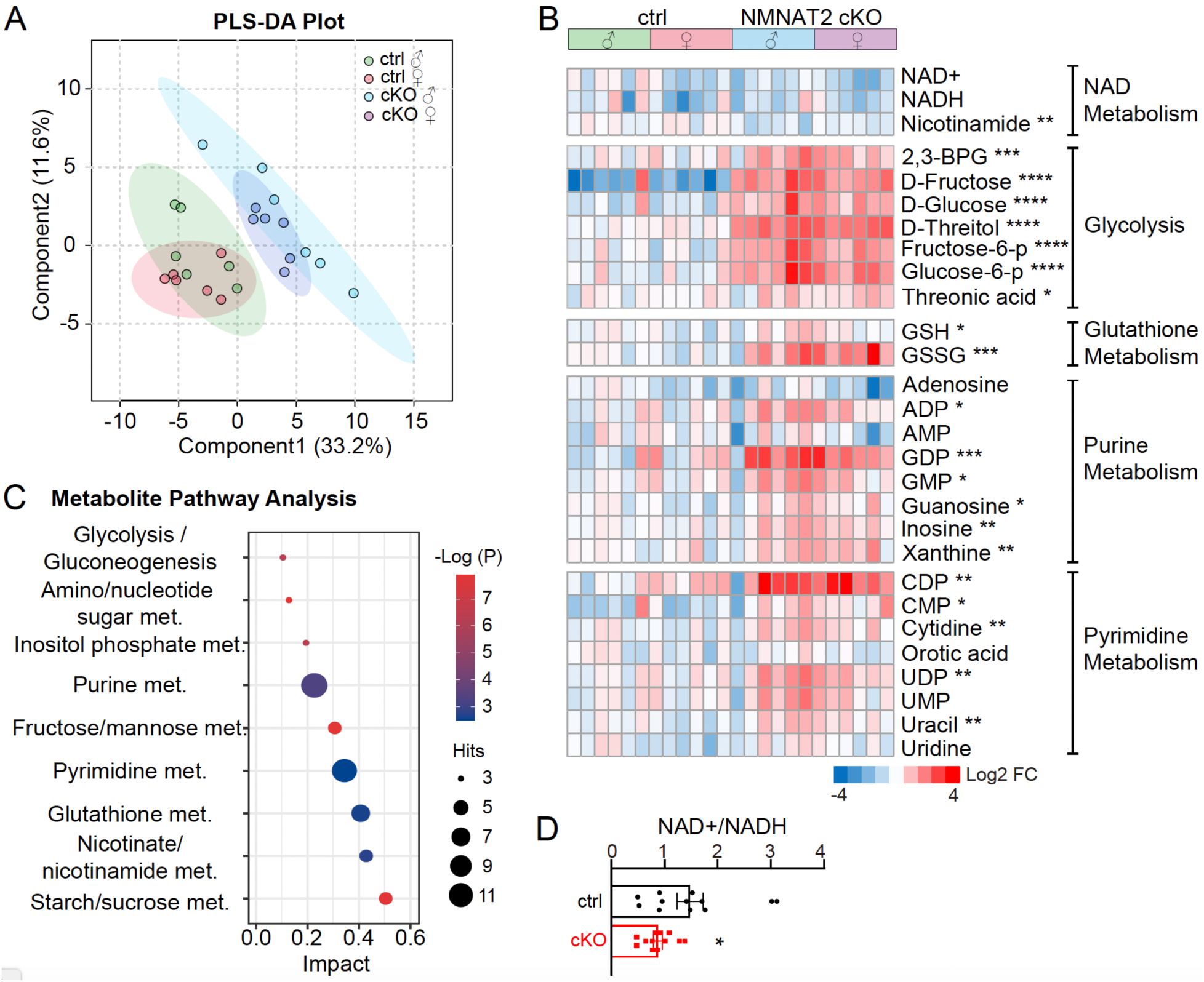
Metabolomic analyses of GC- and LC-MS data reveal differences in several metabolic pathways after NMNAT2 loss in cortical glutamatergic neurons. **(A)** Dimensional reduction and discrimination of individual control (ctrl) and NMNAT2 cKO (cKO) cortices metabolomic profiles using Partial Least Squares Discriminant Analysis (PLS-DA). **(B)** Heatmaps of log2-transformed values summarize the top differential metabolites between cKO and control cortices. They are clustered by metabolic pathways annotated to the right. **(C)** The bubble plot illustrates the impacts of metabolic pathways identified through Metabolite Set Enrichment Analysis (MSEA) with differential metabolites found in cKO cortices. (**D**) Summary for NAD^+^/NADH ratios. Sample sizes: n=6 per sex per group. Abbreviations: cKO, cortical glutamatergic neuron-specific deletion of NMNAT2; met., metabolism. *, p<0.05; **, p<0.01; ***, p<0.001; ****, p<0.0001 by Student’s t-test and Mann-Whitney test.

### NMNAT2 loss in glutamatergic neurons disrupted glucose and lipid metabolism

To gain further insights into the metabolic shifts and determine whether the protein abundance of key enzymes in the identified metabolic pathways could explain the metabolic changes caused by NMNAT2 loss, we conducted proteomic profiling of P16-P21 control and NMNAT2 cKO cortices. PLS-DA analysis of the proteomic data revealed distinct proteomic profiles between the control and cKO groups (Fig. 2A). In NMNAT2 cKO cortices, 292 proteins were significantly upregulated, while 504 proteins were significantly downregulated compared to the control (Fig. 2B). KEGG pathway enrichment analyses were conducted separately with the significantly upregulated and downregulated proteins to identify overrepresented pathways altered in NMNAT2 cKO cortices (Fig. 2C-D left panels). Notably, the fatty acid β-oxidation (FAO) pathway emerged as the most significantly upregulated pathway. In contrast, pathways related to protein processing and lipid biosynthetic pathways, such as steroid biosynthesis, were found to be downregulated. STRING protein-protein interaction network cluster analysis, used to identify key hub proteins, revealed that many upregulated proteins were involved in amino acid and sugar metabolism networks. Interestingly, the analysis also highlighted several upregulated proteins associated with FAO and lipid biosynthetic processes (Fig. 2C right panel). For the downregulated proteins in NMNAT2 cKO cortices, STRING identified several proteins within networks related to lipid metabolism, response to lipids, and protein folding (Fig. 2D right panel). These findings suggest that NMNAT2 loss triggers the upregulation of catabolic pathways while suppressing anabolic (biosynthetic) pathways involved in lipid metabolism. The observed proteomic changes in NMNAT2 cKO cortices emphasize profound alterations in fatty acid/lipid metabolism and protein homeostasis following the loss of NMNAT2 in glutamatergic neurons.

**Fig. 2.**
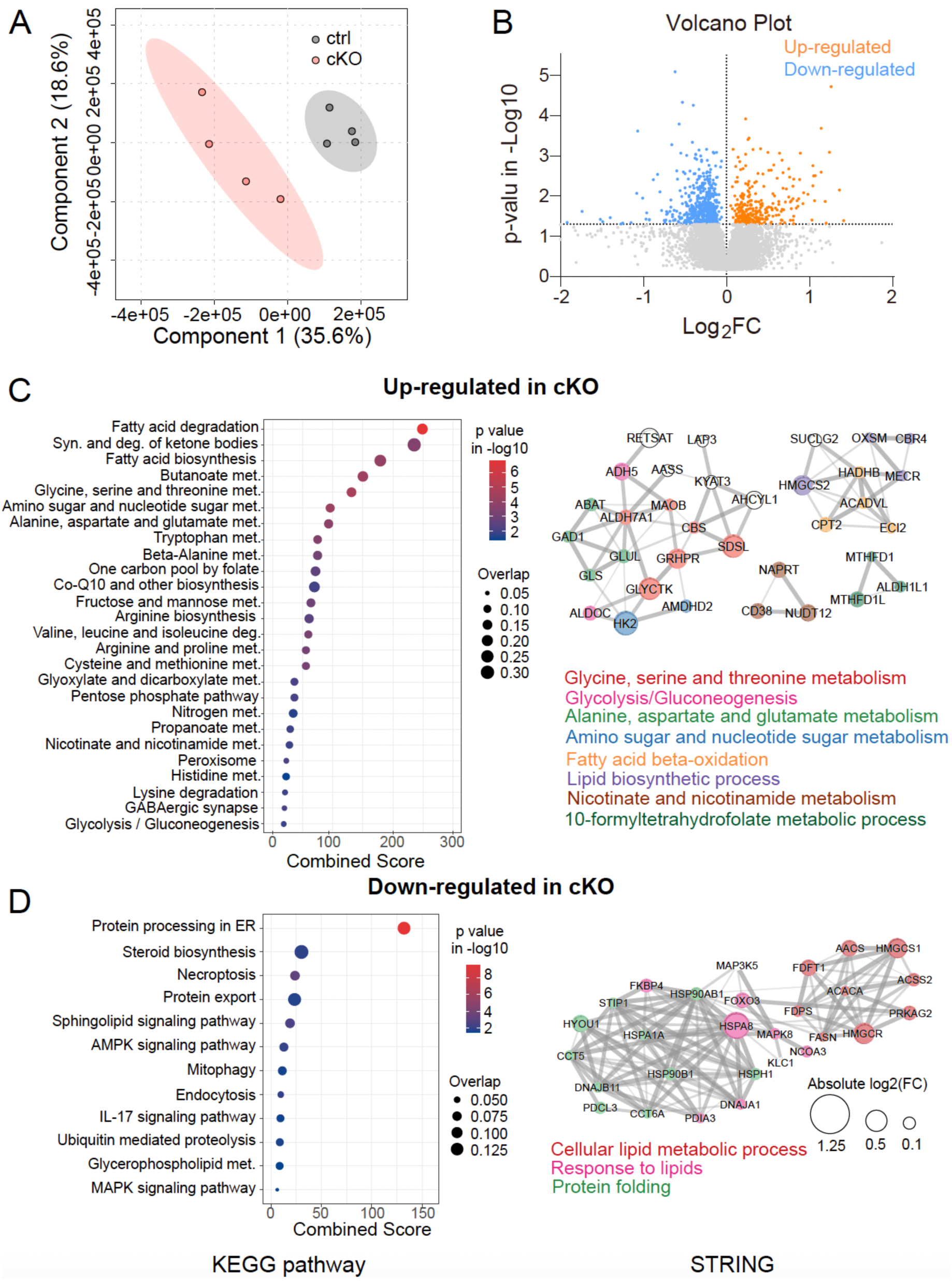
Proteomic profiling of NMNAT2 cKO and control cortices. **(A)** PLS-DA plot shows distinct proteomic profiles of control and cKO cortices. **(B)** A volcano plot shows a large number of significantly up- and down-regulated proteins in cKO cortices (p<0.05). **(C-D)** Summary graphs for KEGG pathway enrichment analysis (left panels) and STRING protein-protein interaction analysis (right panels) with upregulated **(C)** and downregulated proteins **(D)** in cKO cortices. The sizes of bubbles in the bubble plots correspond to the overlap ratio of proteins associated with the indicated pathway, and the x-axis represents the pathway combined score, calculated by multiplying the p-value and z-score of the deviation. The right panels show the key hub proteins identified with STRING, with its edge thickness representing the interaction score and its node size showing the degree of fold change. Sample sizes: n = 4 males per group.

### NMNAT2 loss drives a metabolic shift from glycolysis towards FAO and ketogenesis

To better understand how glucose and lipid metabolic pathways are affected by NMNAT2 loss in neurons, we combined the metabolomic and proteomic data sets to examine the key enzymes and intermediates in each pathway (Figs. 3-4). In NMNAT2 cKO brains, we found substantial increases in several glycolysis intermediates, including glucose, glucose-6-phosphate, fructose-6-phosphate, and 2,3-bisphosphoglyceric acid (Fig. 3A-B). Metabolites in the pentose phosphate pathway (PPP), such as ribulose-5-phosphate and sedoheptulose-7-phosphate, were also increased in the cKO cortices. Proteomic analysis identified increases in glycolytic enzymes, such as hexokinase 2 (HK2) and aldolase (ALDOC), in NMNAT2 KO cortices (Fig. 3C). Such increases in glycolytic enzymes may reflect compensatory responses to the accumulation of upstream glycolytic intermediates. Overall, these results indicate that NMNAT2 loss significantly disrupts glycolysis and the PPP.

**Fig. 3.**
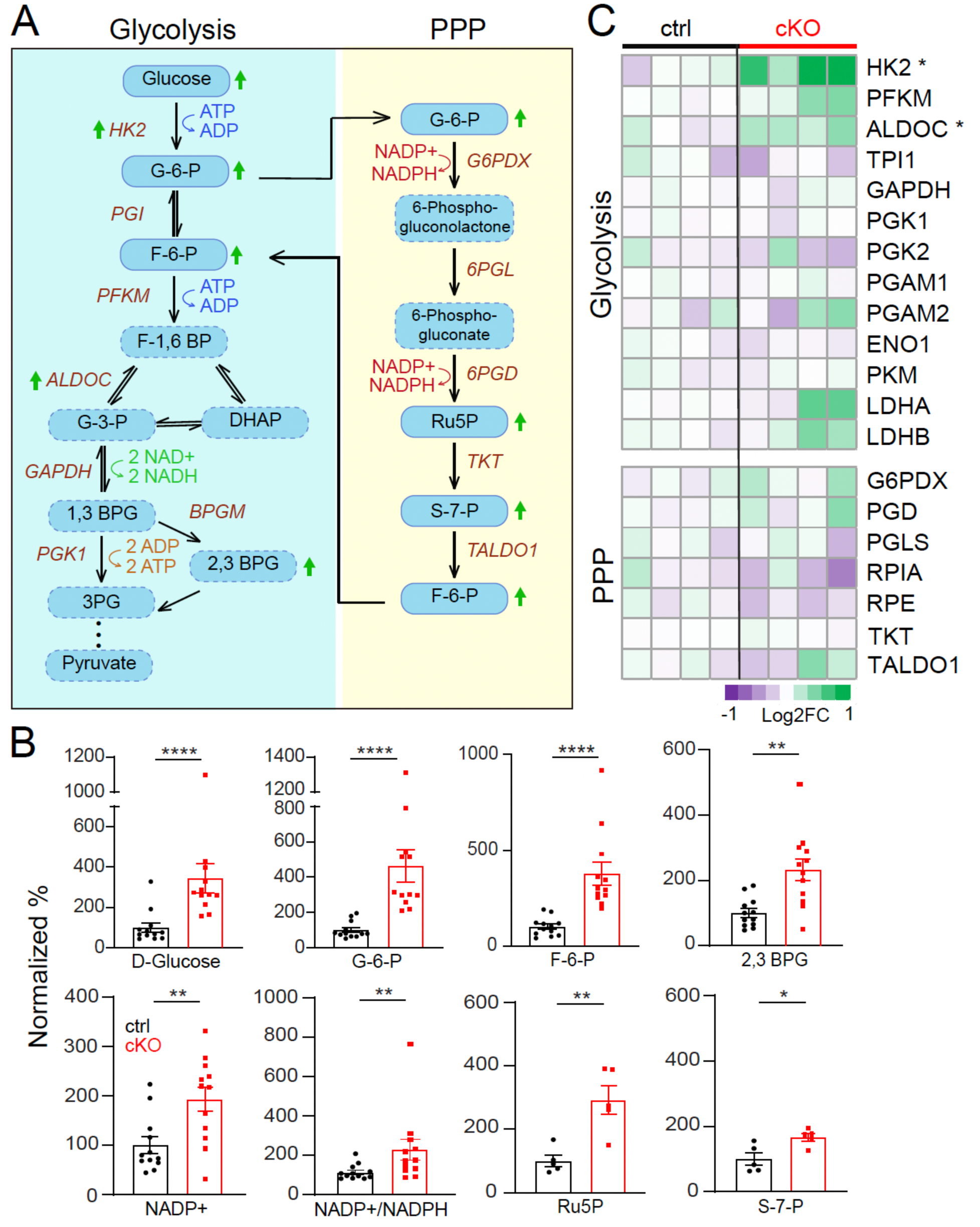
Neuronal NMNAT2 loss results in aberrant glucose metabolism. **(A)** A diagram summarizes key proteins and metabolites involved in glycolysis and the pentose phosphate pathway (PPP). Arrows indicate significant changes in proteins or metabolites. **(B)** Summaries for the normalized abundance of intermediate metabolites in glycolysis and PPP pathways significantly differ between ctrl and cKO cortices (n=6 per sex per group**). (C)** Heatmaps in log2-transformed values summarize the proteomic abundance of enzymes involved in glycolysis and PPP (n=4 per sex per group). *, p<0.05; **, p<0.01; ***, p<0.001; ****, p<0.0001 by Student’s t-test and Mann-Whitney test. Metabolites labeled with a dashed line were not detected in the MS analysis. Abbreviations: 1,3 BPG, 1,3-bisphosphoglycerate; 2,3 BPG, 2,3-bisphosphoglyceric acid; 3PG, 3-phosphoglycerate; ALDOC, fructose-bisphosphate aldolase C; DHAP, dihydroxyacetone phosphate; ENO1, Alpha-enolase; F-1,6 BP, fructose-1,6-bisphosphate; F-6-P, fructose-6-phosphate; G-3-P, glyceraldehyde-3-phosphate; G-6-P, glucose-6-phosphate; G6PDX, glucose-6-phosphate 1-dehydrogenase X; GAPDH, glyceraldehyde-3-phosphate dehydrogenase; HK2, hexokinase-2; LDHA, L-lactate dehydrogenase; LDHB, L-lactate dehydrogenase B chain; PFKM, ATP-dependent 6-phosphofructokinase; PGAM1, phosphoglycerate mutase 1; PGAM2, phosphoglycerate mutase 2; PGD, 6-phosphogluconate dehydrogenase; PGK1, phosphoglycerate kinase 1, PGK2, phosphoglycerate kinase 2; PGLS, 6-phosphogluconolactonase, PKM, pyruvate kinase; RPE, ribulose-phosphate 3-epimerase; RPIA, ribose-5-phosphate isomerase; Ru5P, ribulose-5-phosphate; S-7-P, sedoheptulose-7-phosphate; TALDO1, transaldolase; TKT, transketolase; TPI1, triosephosphate isomerase.

**Fig. 4.**
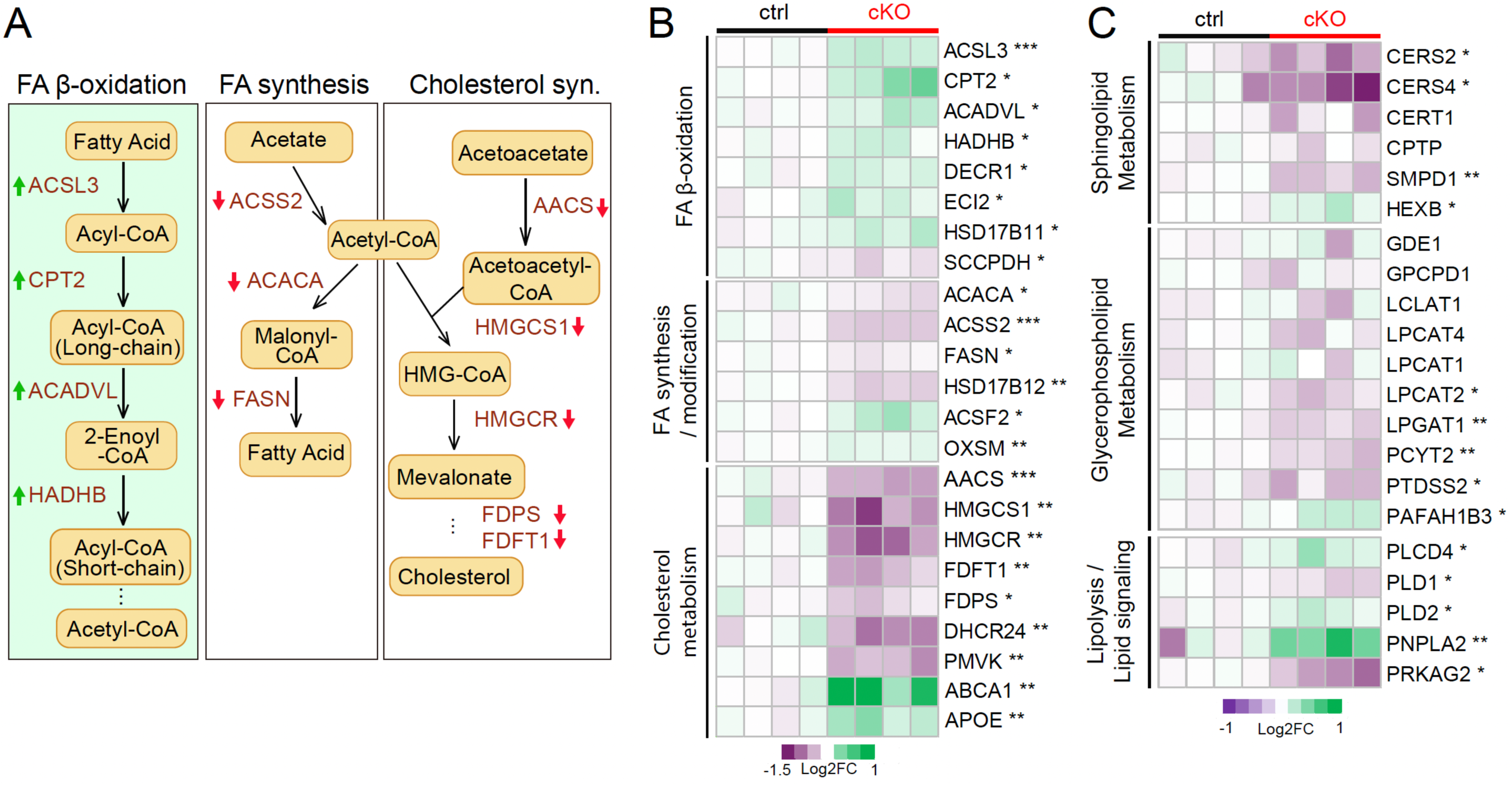
Neuronal NMNAT2 loss results in aberrant lipid metabolism. **(A)** Diagrams summarizing proteins associated with lipolysis and lipogenesis pathways that are differentially expressed in cKO cortices. **(B)** Heatmaps in log2-transformed values summarize the proteomic abundance of enzymes in the FA synthesis/modification, FA β-oxidation, and cholesterol synthesis pathways. Arrows indicate significant changes in proteins or metabolites. **(C)** Heatmap summarizing log2 transformed fold changes of proteomic profiles associated with sphingolipid metabolism, glycerophospholipid metabolism, and lipolysis/lipid signaling. Proteomic sample sizes: n=4 males per group. *, p<0.05; **, p<0.01; ***, p<0.001; ****, p<0.0001 by Student’s t-test and Mann-Whitney test. Abbreviations: AACS, acetoacetyl-CoA synthetase; ABCA1, ATP binding cassette subfamily A member 1; ACACA, acetyl-CoA carboxylase 1; ACADVL, very long-chain specific acyl-CoA dehydrogenase; ACSS2, acyl-CoA synthetase short chain family member 2; ACSF2, medium-chain acyl-CoA ligase; ACSL3, fatty acid CoA ligase; APOE, Apolipoprotein E; CERT1, ceramide transfer protein; CERS2, ceramide synthase 2; CERS4, ceramide synthase 4; CPT2, carnitine palmitoyltransferase 2; CPTP, ceramide-1-phosphate transfer protein; DECR1, 2,4-dienoyl-CoA reductase; DHCR24, delta(24)-sterol reductase; ECI2, enoyl-CoA delta isomerase 2; FASN, fatty acid synthase; FDFT1, squalene synthase; FDPS, farnesyl pyrophosphate synthase; GDE1, glycerophosphodiester phosphodiesterase 1; GPCPD1, glycerophosphocholine phosphodiesterase; HADHB, trifunctional enzyme subunit beta; HEXB, beta-hexosaminidase subunit beta; HMGCR, 3-hydroxy-3-methylglutaryl-coenzyme A reductase; HMGCS1, hydroxymethylglutaryl-CoA synthase; HSD17B11, estradiol 17-beta-dehydrogenase 11; HSD17B12, very-long-chain 3-oxoacyl-CoA reductase; LCLAT1, lysocardiolipin acyltransferase 1; LPCAT1, lysophosphatidylcholine acyltransferase 1; LPCAT2, lysophosphatidylcholine acyltransferase 2; LPCAT4, lysophospholipid acyltransferase; LPGAT1, acyl-CoA:lysophosphatidylglycerol acyltransferase 1; OXSM, 3-oxoacyl-(acyl-carrier-protein) synthase; PAFAH1B3, Platelet-activating factor acetylhydrolase IB subunit alpha1; PCYT2, ethanolamine-phosphate cytidylyltransferase; PLCD4, 1-phosphatidylinositol 4,5-bisphosphate phosphodiesterase delta-4; PLD1, phospholipase D1; PLD2, phospholipase D2; PMVK, phosphomevalonate kinase; PNPLA2, patatin-like phospholipase domain-containing protein 2; PRKAG2, 5’-AMP-activated protein kinase subunit gamma-2; PTDSS2, phosphatidylserine synthase 2; SCCPDH, saccharopine dehydrogenase-like oxidoreductase; SMPD1, sphingomyelin phosphodiesterase.

Regarding lipid metabolism, we found significant upregulation of key enzymes involved in FAO in NMNAT2 cKO tissue (Fig. 4), such as carnitine palmitoyltransferase 2 (CPT2) and acyl-CoA dehydrogenase (ACADVL). The ketogenesis enzymes, such as hydroxymethylglutaryl-CoA synthase (HMGCS2) and D-beta-hydroxybutyrate dehydrogenase (BDH1) are upregulated in cKO cortices (Sup Fig. 1A-B). Together with the increased levels of 3-hydroxybutyric acid (BHB), a ketone body, our data suggest that ketogenesis is enhanced in NMNAT2 cKO cortices (Sup Fig. 1C). Concurrently, several proteins involved in lipogenesis and cholesterol synthesis pathways, including acetyl-CoA carboxylase alpha (ACACA) and acetoacetyl-CoA synthetase (AACS), were significantly downregulated (Fig. 4A-B). Additionally, proteins involved in lipid transport, such as ATP binding cassette subfamily A member 1 (ABCA1) and Apolipoprotein E (APOE), were upregulated in cKO cortices. In summary, our data indicate a metabolic shift in NMNAT2 cKO cortices towards FAO and ketogenesis, while glycolysis and lipolysis were significantly reduced.

### NMNAT2 loss in neurons triggers massive lipid catabolism and complete SARM1 deletion restores its lipid levels

The prominent changes in proteins involved in fatty acid metabolism observed in NMNAT2 cKO cortices (Fig. 4) prompted us to conduct a quantitative lipidomic analysis. Principal component analysis (PCA) showed clear separations between P16-21 cKO and control cortices (Sup Fig. 2A). Strikingly, we found a substantial reduction in total lipid abundance in cKO cortices compared to control (Fig. 5A). Among the 1,254 lipid species identified, 408 were significantly altered in the cKO cortices, with 382 being downregulated (Sup Fig. 2B). Several subgroups of sphingolipids and glycerophospholipids were downregulated in the cKO cortices (Fig. 5B,D; Sup. Table 1). Only a few lipid subgroups, such as bile acids and ceramide phosphates, are elevated in cKO cortices. Notably, these lipidomic alterations correlate with reduced expression of key enzymes involved in glycerophospholipid and sphingolipid metabolism in cKO cortices (Fig. 4C). For instance, ceramide synthases (CERS2, CERS4) and sphingomyelin phosphodiesterase (SMPD1) are downregulated, along with Lysophosphatidylglycerol Acyltransferase 1 (LPGAT1), Lysophosphatidylcholine Acyltransferase 2 (LPCAT2), Phosphatidylserine Synthase 2 (PSS2), and Phospholipase D1 (PLD1).

**Fig. 5.**
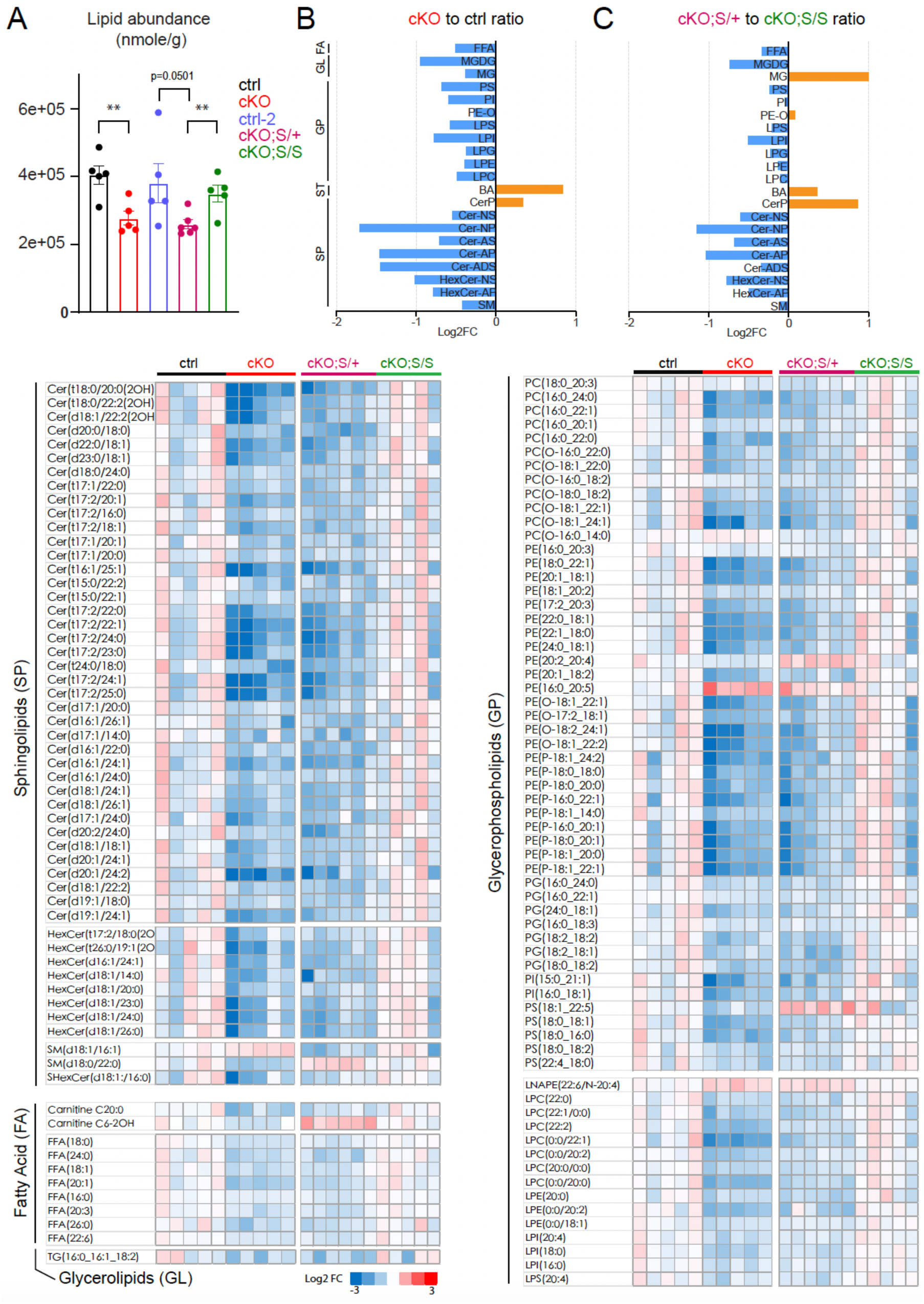
NMNAT2 loss results in a significant reduction in lipid abundance, and complete SARM1 deletion restores the lipid levels. **(A)** Summary for the lipid abundance for control, cKO, control-2, cKO;S^null/+^ (labeled in panels as cKOS/+), and cKO;S^null/null^ (labeled in panels as cKO;S/S) cortices. *, p<0.05; **, p<0.01; ***, p<0.001; ****, p<0.0001 by multiple t-test. **(B-C)** Summary of log2 transformed fold changes for the most highly differentially expressed lipid subclasses. **(D)** Heatmaps highlight the top-ranked differences in lipid content between groups. Sample sizes: n = 5-6 for ctrl-2, cKO, cKO;S^null/+^, and cKO;S^null/null^ males. Abbreviations: cKO;S/+, cKO;S^null^/+; cKO;S/S, cKO;S^null^/S^null^; BA, Bile Acid; Cer-ADS, Ceramide Alpha-Hydroxy Fatty Acid-Dihydrosphingosine; Cer-AP, Ceramide Alpha-Hydroxy Fatty Acid-Phytosphingosine; Cer-AS, Ceramide Alpha-Hydroxy Fatty Acid-Sphingosine; Cer-NP, Ceramide Non-Hydroxy Fatty Acid-Phytosphingosine; Cer-NS, Ceramide Non-Hydroxy Fatty Acid-Sphingosine; CerP, Ceramide-1-phosphate; FA, Fatty Acids; FFA, Free Fatty Acid; GL, Glycerolipids; GP, Glycerophospholipids; HexCer-AP, Hexosylceramide Alpha-Hydroxy Fatty Acid-Phytosphingosine; HexCer-NS, Hexosylceramide Non-Hydroxy Fatty Acid-Sphingosine; LPC, Lysophosphatidylcholine; LPE, Lysophosphatidylethanolamine; LPG, Lysophosphatidylglycerol; LPI, Lysophosphatidylinositol; LPS, Lysophosphatidylserine; MG, Monoacylglycerol; MGDG, Monogalactosyldiacylglycerol; PE-O, Ether-linked Phosphatidylethanolamine; PI, Phosphatidylinositol; PS, Phosphatidylserine; SM, Sphingomyelin; SP, Sphingolipids; ST, Sterol lipids.

We and others have shown that genetically deleting SARM1 protects against the axonopathy caused by NMNAT2 loss ^13,25,28,29^. Here, we aim to investigate whether deleting SARM1 can mitigate the metabolic reprogramming caused by neuronal NMNAT2 loss and subsequently restore brain health. NMNAT2 cKO mice were crossed with SARM1 knockout mice (S^null/null^) to generate NMNAT2 cKO mice in heterozygous (cKO;S^null/+^) or homozygous (cKO;S^null/null^) SARM1-null backgrounds. Cortices from cKO;S^null/+^, cKO;S^null/null^, and littermate control mice from this breeding strategy (named as control-2 to be distinguished from cKO littermate control despite the same genotype as Nex^cre/+^;NMNAT2^f/+^;S^+/+^) were collected for quantitative lipidomic analysis. We have found that the brains of control-2 and cKO;S^null^/S^null^ mice appear morphologically normal, whereas degenerative morphologies are observed in the brains of cKO and cKO;S^null/+^ mice ^13^. The lipidomic analysis shows that the total lipid abundances in cKO;S^null/+^ cortices are significantly reduced, similar to the levels detected with cKO cortices (Fig. 5A). In contrast, control-2 and cKO;S^null/null^ cortices show normal lipid levels comparable to cKO littermate control.

In cKO;S^null/+^ cortices, 234 lipids were significantly altered, with 204 downregulated compared to cKO;S^null/null^ cortices (Sup.Fig.2C-D). Several subgroups of sphingolipids and glycerophospholipids were downregulated, while bile acids and ceramide phosphates were upregulated in the cKO;S^null/+^ cortices compared to cKO;S^null/null^ cortices (Fig. 5C,D). The most similar lipid changes in abundance for cKO and cKO;S^null/+^ compared to control and cKO;S^null/null^, respectively, are bile acids and lipids within the sphingolipid and sterol lipid subclasses (Sup. Table 1). KEGG pathway analysis with altered lipid species reveals the common changes in many pathways, including glycerophospholipid metabolism, sphingolipid metabolism, retrograde endocannabinoid signaling, insulin resistance, necroptosis, and autophagy (Sup. Table 2). In summary, our analysis reveals that, upon the complete deletion of SARM1, many lipid alterations in cKO cortices become normalized.

### NMNAT2 loss alters nucleotide and glutathione metabolism, while SARM1 deletion ameliorates these changes

LC-MS analysis targeting redox metabolites and nucleotides was conducted with the cortices prepared from control-2, cKO;S^null/+^, and cKO;S^null/null^ mice. Surprisingly, no difference in NAD^+^/NADH ratios between cKO;S^null/+^ and control-2 or cKO;S^null/null^ cortices (ratios: control-2, 1.12±0.15; cKO;S^null/+^, 0.91 ± 0.07; cKO;S^null/null^, 0.84±0.10) were observed. However, we found significant changes in several metabolites within the purine and pyrimidine salvage pathways in cKO;S^null/+^ compared to control-2 or cKO;S^null/null^ cortices (Fig. 6A-B). Similar changes in purine/pyrimidine metabolite abundance were also identified in cKO cortices compared to controls. NAD^+^ regulates nucleotide metabolism ^40^, and research shows that NAD^+^ impairment alters the purine and pyrimidine metabolic pathways ^42^. Thus, despite the normalization of NAD^+^/NADH ratios upon the deletion of one copy of SARM1, there are likely to be significant deficits in local NAD^+^ homeostasis to perturb the purine and pyrimidine pathways in cKO;S^null/+^, while complete SARM1 deletion ameliorates the deficits.

**Fig. 6.**
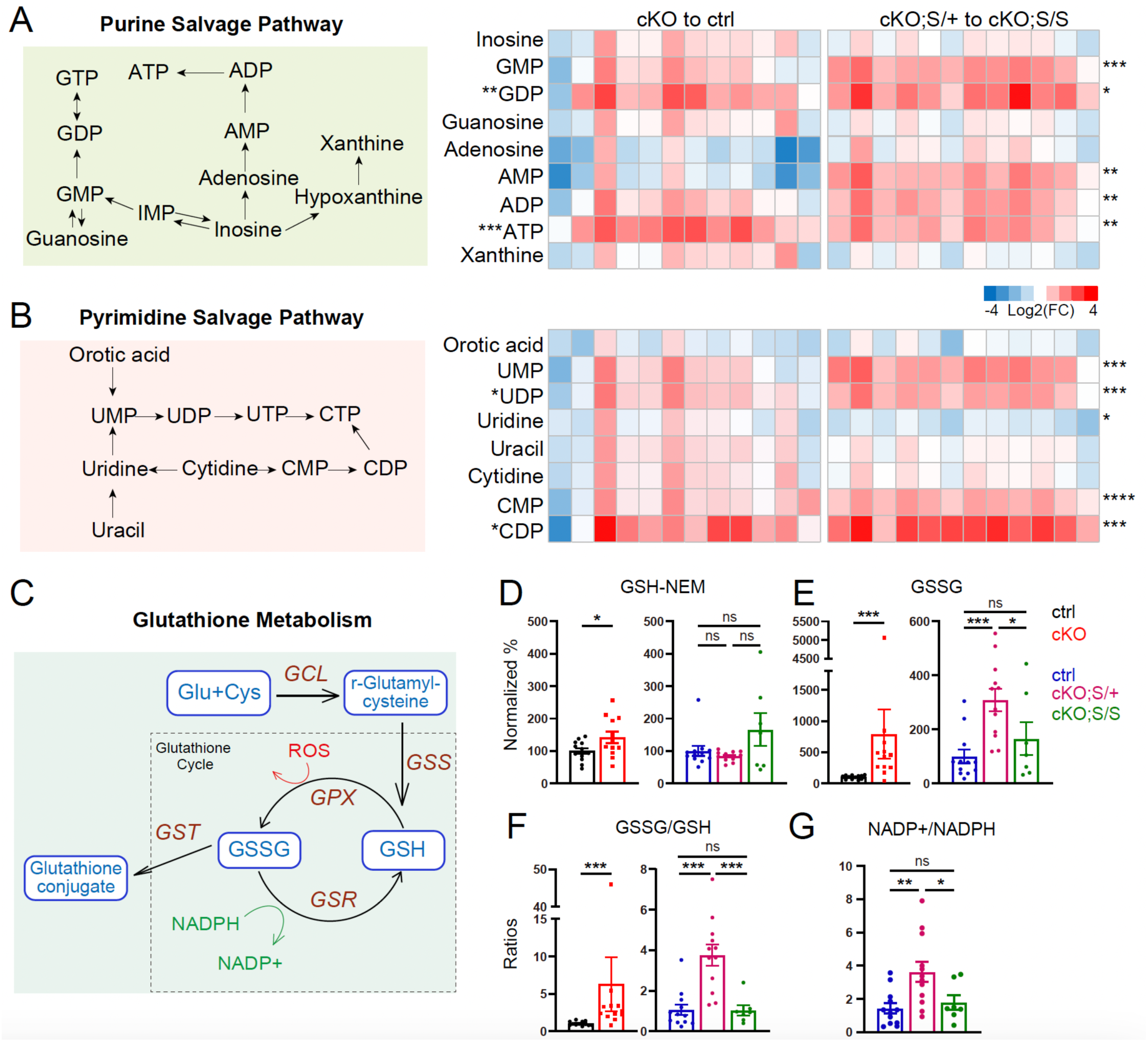
(**A**-**B**) NMNAT2 loss disrupts the nucleotide salvage pathways and complete SARM1 deletion restores the pathways. **(A)** Schematic representation of the purine salvage pathway and heatmaps summarizing fold changes in intermediate metabolite levels for cKO to ctrl and cKO;S^null/+^ to cKO;S^null/null^. **(B)** A diagram of the pyrimidine salvage pathway and heatmaps summarizing fold changes in intermediate metabolites. (**C**) Glutathione metabolism diagram. (**D-E**) Summary of normalized percentages of GSH and GSSG. (**F-G**) Summary of GSSG/GSH and NADP+/NADPH ratios. Sample sizes: NMNAT2 cKO group: n=6 per sex per group; NMNAT2-SARM1 double transgenic group: ctrl-2, n=6 per sex; cKO;S^null/+^, n=6 per sex; cKO;S^null/null^, n=3 males, 4 females. *, p<0.05; **, p<0.01; ***, p<0.001; ****, p<0.0001 by Student’s t-test and Kruskal-Wallis test.

Our targeted LC-MS analysis also revealed elevated ratios of oxidized glutathione (GSSG) to reduced glutathione (GSH) in cKO;S^null/+^ compared to control-2 or cKO;S^null/null^ cortices (Fig. 6C-F). Significant increases in GSSG/GSH (Fig. 6F) ratios were also observed in cKO cortices compared to controls. Glutathione is the most abundant non-protein thiol in mammalian cells, and it acts as a vital reactive oxygen species (ROS) scavenger crucial for cellular defense against oxidative stress ^43–46^. The conversion of GSSG back to GSH, essential for maintaining redox balance, is facilitated by NADPH through the catalysis of glutathione reductase (GR) ^45,47^. Interestingly, NADP^+^/NADPH ratios are elevated in cKO and cKO;S^null/+^ cortices compared to control, control-2, or cKO;S^null/null^ cortices, respectively (Fig. 3B, Fig. 6G). The increased NADP+/NADPH ratios may impair catalysis by GR, consequently increasing GSSG/GSH ratios.

### Loss of neuronal NMNAT2 triggers inflammatory responses and motor behavior deficits, which are mitigated by SARM1 KO

Our proteomic data showed higher abundances of GFAP (a marker for reactive astrocytes) ^48,49^, CD68 (a microglia marker ^50^), and other glia-specific proteins (Sup. Table 3) in NMNAT2 cKO cortices. Additionally, several complement system components and IL-18, a pro-inflammatory cytokine, are also increased in cKO cortices. These proteomic data indicate inflammatory responses in NMNAT2 cKO brains. Inflammation and axonal loss are hallmarks of many neurodegenerative diseases ^51,52^. To examine when NMNAT2 loss results in neurodegenerative phenotypes, immunostainings for GFAP, IBA1 (a microglia marker ^53^), and neurofilament-M (NF-M; a marker for axons) were conducted in coronal brain sections prepared from P5, P16/21, and P90 control and cKO mice. Similar to our previous findings, NF-M+ axon densities at P5 are similar in cKO and control groups in the white matter and striatum, where long-range cortical axons traverse the basal ganglia (Fig. 7A and ^13^). However, after P16/21, NF-M+ axons are significantly reduced. Elevated GFAP signals peaked at P16/21 in the hippocampus, fimbria, white matter, and striatum of cKO mice (Fig. 7B1-B2 and Sup. Fig. 3A-B). GFAP+ astrocytes in cKO brains exhibited bushy morphologies with thickened processes (Fig. 7B2 and Sup. Fig. 3C), a characteristic feature of reactive astrocytes during inflammation ^54^. Additionally, we observed a significant increase in the densities of IBA1+ microglia in the cKO hippocampus and in the cKO striatum from P5 to P90 (Fig. 7B3 and Sup. Fig. 4A-B). Microglia in cKO brains often had large somas with thicker but shorter processes (Fig. 7B3 and Sup. Fig. 4C), which is a characteristic of reactive microglia ^55^.

**Fig. 7.**
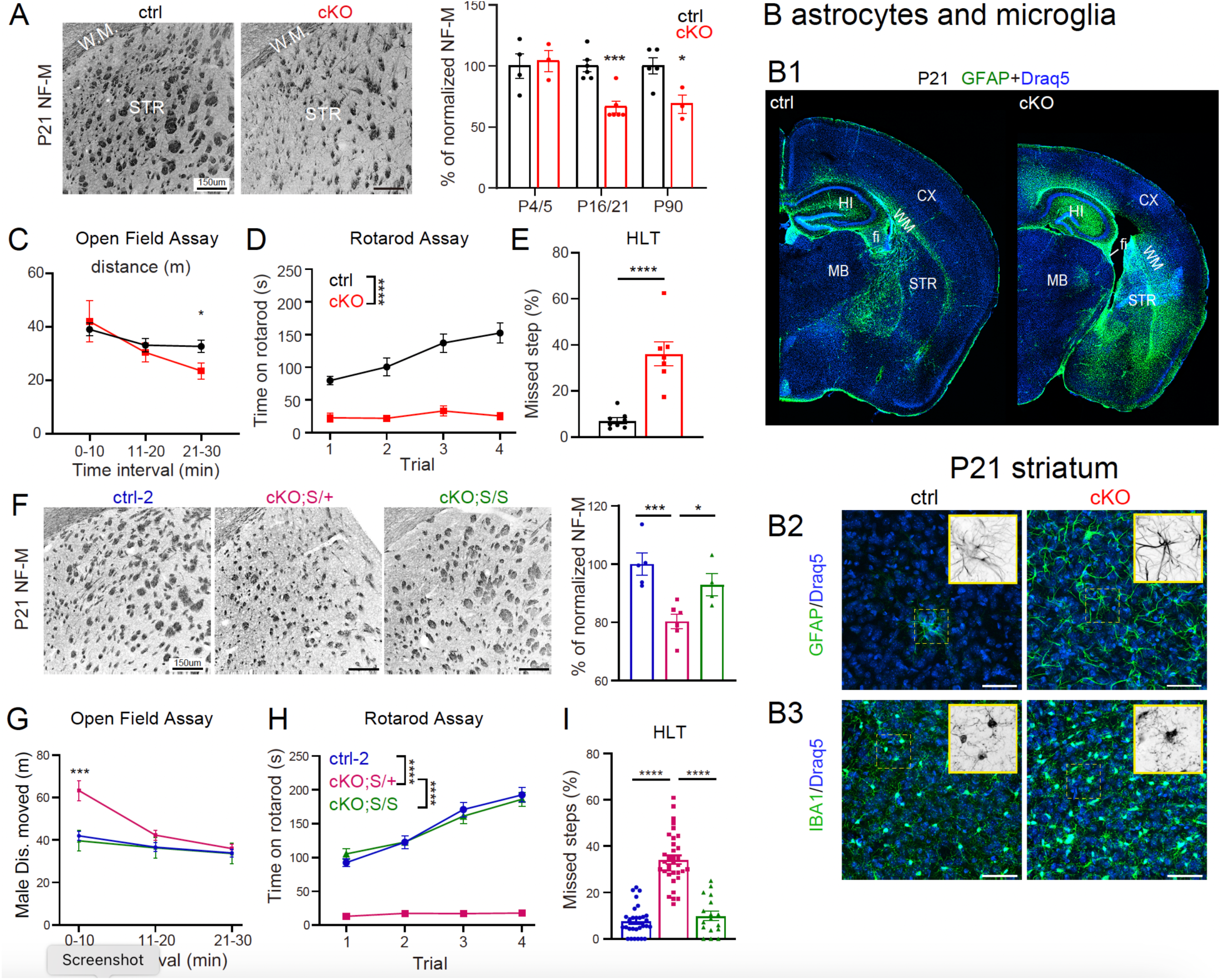
Deleting NMNAT2 in glutamatergic neurons leads to axonal loss, inflammation, and motor deficits, while complete SARM1 deletion improves the neurodegeneration phenotypes. **(A)** Representative NF-M immunostaining images reveal axonal tracts in the striatum (left panels). Bar graphs (right panel) summarize normalized NFM pixel densities detected in the striatum at P4/5 (ctrl, n=4; cKO n=3), P16/21 (ctrl, n=6; cKO n=6), and P90 (ctrl, n=5; cKO, n=3). **(B1-B4)** Stitched confocal GFAP staining images (Draq 5 labels nuclei) of coronal sections from ctrl and cKO brains (**B1**). **(B2)** Representative GFAP staining images from P21 ctrl and cKO striatum. **(B3)** Representative IBA1 staining images. **(C)** Summary for total distance traveled by ctrl and cKO mice in the open field arena during the 30-minute session (ctrl, n=11; cKO n=8). **(D)** Summary of time on rotarod of ctrl and cKO mice during the accelerating rotarod test (ctrl, n=9; cKO, n=8). **(E)** Summary for the percentage of missed steps during horizontal ladder walking test (HLT) (ctrl, n=8; cKO, n=7). **(F)** Representative NF-M immunostaining images in the striatal area of P21 control-3 (ctrl-3; see Methods for details) cKO;S^null/+^, and cKO;S^null/null^ brains. **(G-I)** Summaries for OFA (**G**; ctrl-2, n=15; cKO;S^null/+^, n=15; cKO;S^null/null^, n=8), rotarod (**H**; ctrl-2, n=30; cKO;S^null/+^, n=33; cKO;S^null/null^, n=18) and HLT tests (I; ctrl-2, n=29; cKO;S^null/+^, n=34; cKO;S^null/null^, n=17). *, p<0.05; **, p<0.01; ***, p<0.001; ****, p<0.0001 by ANOVA, Student’s t-test, or Mann-Whitney test. Abbreviations: CX, cortex; HI, hippocampus; WM, white matter; DG, dentate gyrus; STR, striatum; fi, fimbria; MB, midbrain.

Next, we examined if adult NMNAT2 cKO mice exhibit motor behavior deficits in the open-field assay (OFA), the accelerating rotarod test, and the horizontal ladder walk test (Fig. 7C-E). The cKO mice exhibited relatively normal locomotion for the first 20 minutes in the OFA (Fig. 7C). However, they had significant difficulties in staying on the accelerating rotarod, as indicated by markedly reduced time on the rotarod (Fig. 7D). In contrast to control mice, which had very few missed steps while walking on the horizontal ladder, cKO mice missed significantly more steps (Fig. 7E). These motor behavioral impairments noted in cKO mice suggest deficits in their motor coordination.

cKO;S^null/+^ mice also displayed reduced NF-M+ axon densities (Fig. 7F) and increaseed GFAP and IBA1 immunoreactivity in the striatum compared to control-2 and cKO;S^null/null^ mice (Sup. Fig. 5). Excitingly, the complete absence of SARM1 alleviates motor coordination deficits observed in NMNAT2 cKO mice (Fig. 7H-I), while cKO;S^null/+^ mice show hyperlocomotion during the first 10 minutes in OFA (Fig. 7G), severe impairments in the accelerating rotarod (Fig. 7H) and horizontal ladder walk (Fig. 7I) tests. Collectively, the immunostaining and behavioral data suggest that the total loss of SARM1 ameliorates the neuroinflammation, axonopathy, and motor coordination deficits caused by NMNAT2 deficiency.

### Lipid metabolism-related proteome module reversed by SARM1 KO is linked to neurodegenerative phenotype

To gain molecular insight into the rescue offered by complete SARM1 deletion on neurodegeneration caused by loss of NMNAT2 in the cKO cortices, we performed proteomic profiling with control-2, cKO;S^null/+^, and cKO;S^null/null^ cortices. After batch effect reduction and integration with the proteomic datasets from control and cKO cortices shown above (Fig. 2), 6850 proteins commonly detected in all datasets, including control, cKO, control-2, cKO;S^null/+^, and cKO;S^null/null^, were used to conduct the weighted gene correlation network analysis (WGCNA). Proteins exhibiting a similar expression pattern across samples were clustered into individual modules. Among the total 36 modules identified (data not shown), we identified 8 modules that show significant correlations with neurodegenerative phenotypes and NMNAT2 allele dosages (Fig. 8A). Interestingly, only two of these 8 modules exhibited expression patterns that were significantly reversed by SARM1 allele dosage. (Fig. 8A,B). A single-allele deletion of SARM1 readily rescued protein expression in the Steelblue module, whereas complete loss of SARM1 reversed protein changes in the Plum2 module. Interpreted by the STRING PPI network, the Plum2 module is chiefly enriched in cellular lipid metabolic process and protein folding, followed by chemokine activity, and neuronal dense core vesicle transport. AACS, HMGCS1, FDFT1, and PMVK are involved in cholesterol metabolism (Fig. 4 and Sup. Table 3), ACSS2 and FASN mediate fatty acid synthesis, and PCYT2 catalyzes the second step of phosphatidylethanolamine (PE) synthesis. STRING PPI network analysis with proteins in the Steelblue module identifies proteins involved in oxidoreductase activity, cholesterol biosynthesis, and protein processing in the ER. The WGCNA outcome, along with our lipidomic data, shows that SARM1 deletion alleviates lipid metabolic deficits positively associated with the neurodegenerative phenotype and negatively associated with the NMNAT2 allele.

**Fig. 8.**
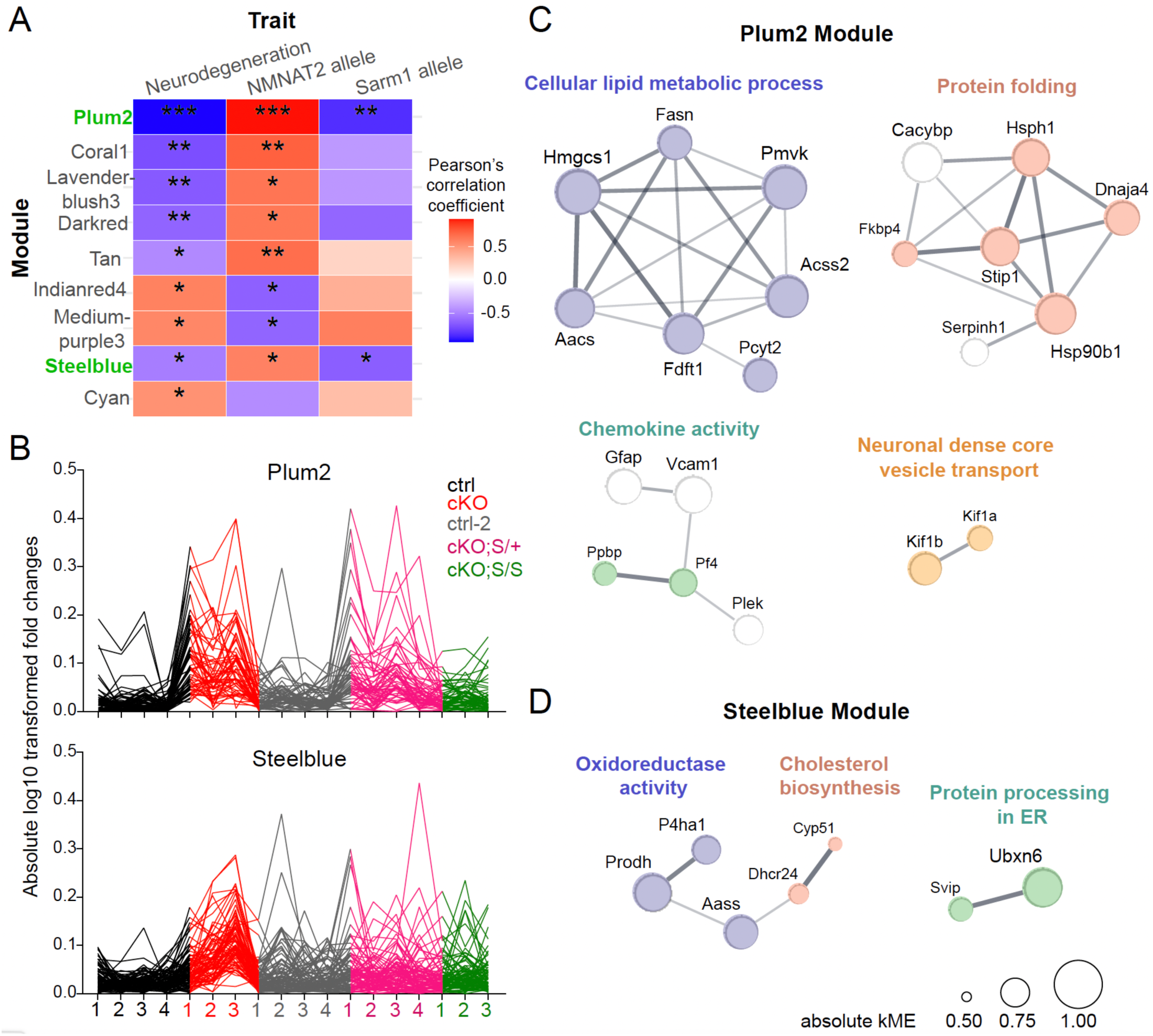
Bioinformatic analysis with WGCNA identified several protein modules that significantly correlated with neurodegenerations and NMNAT2 abundance. SARM1’s presence exerts similar impacts as neurodegeneration on two protein modules. **(A)** Module-trait correlation heat map. Color denotes the Pearson correlation coefficient, p<0.05 *, p<0.01 **, p<0.001 ***. **(B)** Individual protein expression patterns across samples within the Plum2 and Steelblue module. Absolute log10 transformed fold changes were derived from the original protein abundance value before batch-effect reduction. **(C-D)** PPI network of Plum2 (**C**) and Steelblue (**D**) module proteins, with its edge thickness representing the interaction score and its node size showing the eigenprotein-based connectivity (kME), which is the Pearson correlation between the expression profile of a gene and the first principal component of the expression matrix of all proteins within the module (eigenprotein).

## Discussion

Maintaining NAD^+^ homeostasis is critical to support brain energy metabolism and health. NMNAT2 is the most abundant NAD^+^-synthesizing enzyme in cortical neurons ^12^ and is required for neuronal health ^13,14,56^. Its reduction or absence activates SARM1, a NAD^+^ hydrolase, leading to further NAD^+^ depletion and axon degeneration ^21,22,28^. In this study, we used mice lacking NMNAT2 in post-mitotic glutamatergic neurons to determine how NAD^+^ reduction in neurons impacts energy homeostasis in the cerebral cortex. Multi-omics analyses were conducted to determine the proteomic, metabolomic, and lipidomic landscape changes caused by NMNAT2 loss. Notably, we observed that neuronal NMNAT2 loss markedly increases fatty acid catabolism while decreasing lipid synthesis and leads to the accumulation of large amounts of sugars and their derivatives. The complete genetic deletion of SARM1 is required to restore the total lipid abundance and halt the neurodegenerative phenotypes that generally occur in NMNAT2 KO mice. We also found that SARM1 deletion restores altered purine, pyrimidine, and glutathione metabolism in NMNAT2 cKO cortices. These findings not only highlight the essential role of NMNAT2 in maintaining brain energy homeostasis but also reveal the risks that metabolic maladaptation poses to lipid catabolism.

### Impact of NMNAT2 loss on metabolic pathways and the mitigating effects of SARM1 deletion

Our study shows that NMNAT2 loss in glutamatergic neurons triggers significant alterations in several metabolic pathways in the cortex, including nicotinate/nicotinamide, carbohydrate, lipid, nucleotide, and glutathione metabolisms. The observation of a substantial accumulation of sugar and sugar derivatives in NMNAT2 cKO brains was surprising, given the presence of other NMNATs and the specific deletion of NMNAT2 in glutamatergic neurons. The elevated glucose, glucose-6-phosphate, and fructose-6-phosphate levels, together with PPP intermediate metabolites, such as ribulose-5-phosphate and sedoheptulose-7-phosphate (Fig. 3A,C), suggest elevated flux through the PPP in NMNAT2 cKO cortices.

The top pathways dysregulated upon NMNAT2 loss identified by proteomic analysis are primarily lipid metabolic pathways, including those involved in fatty acid synthesis/degradation/β-oxidation and synthesis/degradation of ketone bodies (Fig. 2C). Our lipidomic analysis revealed a substantial reduction of total lipid levels in NMNAT2 cKO cortices (Fig. 5; Sup. Table 1). Many subgroups of sphingolipids and glycerophospholipids are down-regulated in cKO cortices. Studies show that their metabolic processes are altered in various neurodegenerative diseases conditions ^61,62^. Proteomic data show that NMNAT2 cKO cortices had reduced levels of key lipid synthesis enzymes, such as ACSS2, ACACA, and FASN, while higher levels in key enzymes involved in FAO, such as ACADVL, ACSL3, and CPT2 (Fig. 4) were increased. Together with the dramatic reduction in total lipid abundance, our data suggest that NMNAT2 loss decreases cortical lipogenesis while enhancing cortical lipid catabolism in the cortex. Lipid metabolism is fundamental for maintaining neurotransmission, cellular structure, and signaling ^57–60^. Thus, the perturbed lipid metabolism upon NMNAT2 loss is likely to contribute to many of the phenotypes detected with proteomic analysis, histology, and behavioral assays.

The brain’s high energy demand is primarily met by metabolizing glucose ^63–65^. It has a limited lipid storage capacity and depends on tightly regulated lipid synthesis, turnover, and utilization to maintain homeostasis ^2,33,34,66^. However, brain metabolism is plastic, and the brain can use alternative fuel sources in response to metabolic stresses such as starvation and glycolytic impairment ^31,67–70^. In this regard, ketones such as acetoacetate and β-hydroxybutyrate (BHB) represent an important alternative energy source for the brain. While ketogenesis predominantly occurs in hepatocytes ^70,71^, *in vitro* studies suggest that astrocytes can synthesize ketone bodies due to their ability to oxidize fatty acids ^32,70,72^. Our findings support this hypothesis by revealing that NMNAT2 cKO cortices exhibit an increased abundance of the ketogenesis-related enzymes, HMGCS2 and BDH1, as well as elevated levels of the ketone BHB (Sup. Fig 1), which was recently shown to be metabolized by neurons in a cell-autonomous manner to fuel synaptic transmission ^37^. Thus, our proteomic data suggest an enhanced ketogenesis in NMNAT2 KO cortices, but we cannot rule out an increase of ketones from peripheral sources.

Our proteomic studies found that ABCA1 and APOE abundance are upregulated in NMNAT2 cKO cortices. APOE is the main protein component of lipoprotein particles produced by astrocytes and microglia, and it mediates various aspects of lipid homeostasis in the brain ^73–76^. ABCA1 is the primary contributor of lipids to APOE-lipoproteins in the extracellular space ^77^. Such lipidated APOE-lipoproteins supply neurons with cholesterol and phospholipids from astrocytes, while also shuttling lipid peroxides from neurons to astrocytes for detoxification. Though our lipidomic analysis did not detect changes in cholesterol levels, our proteomic analysis revealed reductions in several cholesterol-synthesizing enzymes, such as AACS and HMGCS, in NMNAT2 cKO cortices (Fig. 4).

Neurodegenerative phenotypes in NMNAT2 cKO mice are mitigated by genetic SARM1 deletion. Does the absence of SARM1 activation prevent the reduction in NAD levels and thus avoid the widespread metabolic reprogramming? We found that deleting one copy of SARM1 is sufficient to restore the cortical NAD^+^/NADH ratios. However, the reduced cortical lipid levels in the cortices and neurodegenerative phenotypes are similar between cKO;S^null/+^ and NMNAT2 cKO mice (Figs. 5,7). It is possible that our metabolomic analysis with whole cortices failed to detect localized NAD^+^ metabolism deficits in cKO;S^null/+^ brains and localized NAD^+^ metabolism deficits may be sufficient to cause the maladapted metabolic plasticity. Eliminating two copies of SARM1 in NMNAT cKO mice significantly restores lipid, purine/pyrimidine, and glutathione levels (Fig. 6). Regarding lipid metabolism, the most notable reversal upon total SARM1 deletion is the abundance of lipid species in the sphingolipid and sterol lipid subgroups, as well as bile acids. Congruently, WCGNA analysis of proteomic profiling revealed a protein module that associates with cholesterol, fatty acid, and PE synthesis pathways is negatively correlated to NMNAT2 expression and positively correlated to neurodegenerative phenotypes and SARM1^wt^ alleles (Fig. 8).

### Reducing neuronal NAD^+^ levels activates glia

The metabolism of glia and neurons is extensively coupled ^36,78^. Astrocytes are positioned between neurons and the vasculature to control energy fluxes and nutrient supplies required for neuronal function ^79^. Astrocytes become “reactive” upon metabolic changes in glucose and lipid metabolism ^80^ , while microglia are activated by increases in extracellular purines, pyrimidines, or cytokines ^81,82^. Our proteomic analysis finds that neuronal NMNAT2 loss elevates inflammatory responses, evident by significant increases of reactive glia markers (Sup. Table 3), pro-inflammatory cytokines, and several components found in the complement system in the cKO mice ^83,84^. Histological evaluations of GFAP and IBA1 immunostaining also found inflammatory changes in cKO brains. These are particularly prominent in white matter and in the striatum, which is enriched with long-range axons projecting from glutamatergic neurons (Fig. 7B). This observation suggests that signals originating from the axons of NMNAT2 KO glutamatergic neurons activate neighboring astrocytes and microglia.

Given that astrocytes are the primary brain cells capable of efficiently oxidizing medium- and long-chain fatty acids for energy ^2,85–88^, the upregulation of FAO enzymes and BHB (Sup. Fig. 1) suggests the possibility that astrocytes in NMNAT2 cKO cortices are oxidizing fatty acids to supply neurons with ketone bodies. This is consistent with previous studies showing astrocytes upregulate their FAO and ketogenesis to sustain neuronal energy demands ^65,89^. ACSF2, HMGCR, and PLD2, the upregulated enzymes involved in lipid metabolism in cKO cortices, are predominantly expressed in astrocytes ^90,91^. It remains to be determined whether there is a shift from glycolysis to fatty acid catabolism in astrocytes, which subsequently increases sugars and sugar derivatives in cKO cortices.

An increased abundance of HK2, but not HK1, in NMNAT2 cKO cortices, is a notable finding. HK2 is primarily expressed in microglia and is required for microglial glycolytic flux and energy production ^92,93^. Since microglial activation is often accompanied by elevated glycolysis, increased HK2 expression is believed to support inflammatory responses by sustaining ATP production independently of OXPHOS ^94^. We observe that HK2 is upregulated in parallel to microglia activation (Fig.3 and Sup. Table 3), which is evidence of a glial response to signals from NMNAT2 KO neurons. Interestingly, HK2 antagonism slows neurodegeneration in 5XFAD mice, a commonly used AD mouse model ^95^. ACSS2 in the FA synthesis pathway is downregulated in NMNAT2 cKO cortices (Fig. 4; Sup. Table 3). It has recently been shown that ACSS2 is predominantly expressed in oligodendrocytes ^96^ , and its upregulation results in acetyl-CoA increases for de novo lipogenesis and histone acetylation ^97^. Collectively, our data suggest that NMNAT2 loss in glutamatergic neurons induces profound metabolic reprogramming not only in neurons but also in astrocytes, microglia, and oligodendrocytes.

Impaired glycolysis or altered FAO ^33,66,98^ can induce oxidative stress to activate microglia and astrocytes, subsequently triggering neuroinflammatory responses ^99,100^. In other words, the metabolic plasticity can result in a vicious cycle by triggering chronic inflammation that worsens neuronal damage ^101,102^. Our metabolomic analysis found elevated levels of 3-nitrotyrosine, a marker of oxidative stress ^103^, in cKO cortices (normalized %, Ctrl: 100, cKO: 126.6; Difference: 26.59 ± 8.23, p < 0.05). Could the reactive astrocytes and microglia in NMNAT2 cKO brains contribute to the maladaptive metabolic plasticity? Inflammatory responses with increased astrocyte reactivity and microglia numbers have been observed in many neurodegenerative conditions ^104,105^. Future studies to understand how the disruption of neuronal NAD homeostasis triggers inflammatory responses will offer essential insights for therapeutic interventions designed to reduce inflammation-related harm.

In summary, our data provide strong support that when glucose metabolism in neurons is disrupted by the loss of NMNAT2, the brain increases FAO and ketogenesis to meet its energy demands. Such a shift disrupts the normal lipid metabolism that is critical for neuronal function and healthy neuron-glia communications. SARM1 is required for the FAO that accompanies the loss of NMNAT2. Confocal imaging and biochemistry studies suggest that SARM1 is localized in the mitochondria ^106,107^, where FAO occurs. Thus, future studies will be important to elucidate how SARM1 activity affects FAO. Identifying methods to modify energy metabolism in mitochondria may provide an opportunity to prevent harm from metabolic maladaptation, suppress inflammatory responses, and prevent neurodegeneration.

## Acknowledgment

This work was supported by the following grants: National Institutes of Health grants R01NS086794 (HCL) and P30DA056410 (HCL). We thank Scott Barton and Jason Fu for their technical support, and Drs. Ken Mackie, Anna Kalinovsky for their helpful comments. Metabolomics analysis was performed at the University of Utah’s Metabolomics Core, supported by NCRR 1S10OD016232-01, 1S10OD018210-01A1, and 1S10OD021505-01. The Indiana University School of Medicine Proteomics Core performed proteomic mass spectrometry, supported by NIH UL1TR002529 and NCI P30CA082709.

## Materials and Methods

### Animals and genotyping

Cortical glutamatergic neuron-specific NMNAT2 knockout (Nex^cre/+^;NMNAT2^f/f^; cKO) mice were generated by crossing NMNAT2^f/f^ mice ^15^ with NEX-Cre mice (Nex^cre/+^)^39^ as previously described ^13^. Nex^cre/+^;NMNAT2^f/+^ littermates were used as the control mice. SARM1 knockout (S^null/null^) mice ^107^ were crossed with Nex^cre/+^;NMNAT2^f/+^ mice to generate cKO;S^null/+^ and cKO;S^null/null^ transgenic mice. The littermates with Nex^cre/+^;NMNAT2^f/+^ genotype were used as control (control-2) for lipidomic analysis. The littermate with Nex^cre/+^;NMNAT2^f/+^ or Nex^cre/+^;NMNAT2^f/+^;S^null/+^ genotypes were used as controls (control-3) for neurodegenerative phenotyping (no significant difference in the results from immunostaining and behavioral experiments were observed between these two genotypes). Both NMNAT2 cKO and cKO;S^null/+^ mice exhibited an ataxia phenotype, and thus, a 5g DietGel 76A (Clear H_2_O) was given daily upon weaning.

Both male and female mice were used in this study. All mice were housed under standard conditions with ad libitum access to food and water and maintained on a 12-hour light/dark cycle. Animal care and experimental procedures complied with the guidelines of the U.S. Department of Health and Human Services and were approved by the Indiana University Bloomington Institutional Animal Care and Use Committee. The genotyping procedure was conducted as previously described ^13^

### Tissue preparations and immunostaining

Mice with desired genotypes were anesthetized and transcardially perfused with 4% paraformaldehyde (PFA) in phosphate-buffered saline (PBS). Brains were harvested and post-fixed overnight in 4% PFA at 4°C. After rinsing with PBS, the brains were cryopreserved in 30% sucrose in PBS at 4°C for 1–2 days. Cryopreserved brains from P5-P90 were mounted on the cutting platform of a Leica microtome (SM-2000R, Leica) with dry ice and sectioned into free-floating coronal slices at 30–40 μm thickness for subsequent staining procedures.

The immunostaining process was conducted at room temperature unless otherwise noted. Sections were washed three times for 5 minutes each with PBS containing 0.3% Triton (0.3% PBST), then permeabilized with 0.3% PBST for 20 minutes. Following permeabilization, sections were incubated in a blocking solution (3% bovine serum albumin (BSA) in 0.3% PBST) for 2 hours and then incubated overnight at 4°C with primary antibodies diluted in blocking solution. The following antibodies were used: GFAP antibody (DAKO, GA524, RRID:AB_2811722, 1:1000), IBA1 antibody (WAKO, 019-19741, RRID: AB_839504, 1:1000), and Neurofilament-M antibody (Millipore, MAB1621, RRID:AB_94294, 1:1000). The next day, sections were washed three times for 5 minutes each with 0.3% PBST and incubated with secondary antibodies diluted in blocking solution at 4°C for 2 hours. The following secondary antibodies were used: Alexa Fluor 488 goat anti-Rabbit IgG (H+L) antibody (Thermo Fisher, A-11034, RRID:AB_2576217, 1:1000), Alexa Fluor 488 goat anti-Chicken IgG (H+L) antibody (Molecular Probes, A-11039, RRID:AB_142924, 1:1000), Alexa Fluor 488 goat anti-Mouse IgG (H+L) antibody (Molecular Probes, A-11029, RRID:AB_138404, 1:1000), Alexa Fluor 555 goat anti-Rabbit IgG (H+L) antibody (Molecular Probes, A-21429, RRID:AB_2535850, 1:1000). After incubation, sections were washed three more times with 0.3% PBST. To visualize nuclei, Draq5 (Cell Signaling, 4048L, 1:10,000) or DAPI (Invitrogen, D3571, 5 μg/ml) was applied. Finally, sections were mounted using Dako Mounting Medium (Agilent Technologies, CS70330-2) for imaging.

### Behavioral Assays

All behavior tests listed below were conducted under 100-lux white-light illumination. Mice were transferred to the testing room and acclimated for at least 1 hour before tests. Experiments were performed around a similar time of day to align with the animals’ biological rhythms.

#### Horizontal Ladder Test (HLT)

To assess locomotor skills, the horizontal ladder walking test was performed ^108^. Mice were placed on a 100 cm-long horizontal ladder with unevenly spaced metal rungs, elevated 15 cm above the ground. The mouse’s home cage was positioned at the end of the ladder to encourage movement. The camera was placed laterally to record the animal’s movement along the ladder. From the recorded videos, missteps of both the left and right hind limbs were manually counted. The percentages of missing steps for both the left and right limbs relative to the total steps were calculated.

#### Rotarod Test

The rotarod test (Ugo Basile, 47600) was conducted as previously described ^109^. Briefly, an individual mouse was placed on the rotating rod facing away from the direction of rotation, with an initial speed of 4 RPM. Once all mice were in place, the rotation speed was accelerated from 4 RPM to 40 RPM over 300 seconds. The latency to fall and the rotational speed at which the mouse fell were recorded. Each mouse underwent four consecutive trials, with a 5-minute rest period in their home cage between trials.

#### Open Field Assay

Locomotor activity was assessed using the open field assay. Mice were placed in the center of an open field arena (43 cm × 43 cm × 43 cm) enclosed by non-transparent Plexiglas walls and allowed to move freely for 30 minutes. Travel distances and velocity were recorded using EthoVision XT software, with the animal’s head designated as the reference point for tracking. The total distance traveled was used as a measure of general locomotor activity ^110^. The arena was thoroughly cleaned with 70% ethanol between trials.

### Confocal microscopy and quantitative image analysis

Confocal fluorescence images were captured with either a Leica TCS SPE confocal microscope (Leica DM 2500) using either a 10× objective lens (0.3 N.A.) or a 40× objective lens (0.75 N.A.) or a Nikon A1R laser scanning confocal microscope (Nikon A1) equipped with a 10× objective lens (0.5 N.A.) or a 60× oil-immersion objective lens (1.4 N.A.). DAPI/Draq5 immunofluorescence was used to identify comparable anatomical regions across different brain sections. >3 sections per mouse were imaged for both hemispheres with identical imaging parameters for the antibody staining and analyzed in a consistent and comparable manner across all experimental groups.

The imaging setting was chosen to minimize signal saturation across experimental groups. All quantifications were conducted blind to genotype information. DAPI or Draq5 immunofluorescence was used as a reference to draw the contours of regions of interest.

Images were analyzed using Fiji (ImageJ with updated plug-ins). IBA1-positive cell densities were determined by manually marking positive cells using the ImageJ Cell Counter function. For GFAP-positive astrocyte quantification, images were thresholded using the top 3.7–4.5% tail of total pixels, followed by “Analyze Particles” to calculate pixel percentages. Image acquisition and quantification were performed with 0.25mm^2^ area in the dorsal striatum and 0.075mm^2^ in hippocampal CA1.

For NFM-positive axonal tracts passing through the dorsal striatum, only coronal slices from the middle rostral-caudal axis were selected for quantification. Areas occupied by blood vessels were excluded. The vessel-subtracted dorsal striatal regions were then processed using the “Threshold” function (top 8–11% tail of total pixels). The “Analyze Particles” function was then applied to determine the percentage of pixels representing NFM-positive axons.

### Brain tissue preparation for metabolomic proteomic and lipidomic MS analysis

All the Brain tissues were collected when mice reached postnatal day 16–21. The somatosensory cortical region was rapidly dissected immediately after euthanasia. All procedures were performed on ice to preserve tissue integrity. The dissected tissues were then snap-frozen in a dry ice/ethanol bath and stored at -80°C for subsequent metabolomic and proteomic mass spectrometry analysis.

### KEGG pathway and PPI network analysis for proteomic data

Significantly up- or downregulated proteins (p<0.05) were used for KEGG pathway enrichment analysis through EnrichR. Irrelevant terms, such as “Malaria” and “Salmonella infection”, were removed, and the top 25 up- and top 15 down-regulated pathways are shown. Similarly, significantly up- or downregulated proteins were used for protein-protein interaction (PPI) network construction on the STRING website, using a medium confidence (0.4) interaction score filter. The network edge thickness represents interaction confidence based on experiments, databases, and co-expression evidence. DBSCAN clustering with a network-specific epsilon parameter (network radius times 4) was applied. The clusters with at least 3 members were selected and exported to Cytoscape software for further KEGG pathway enrichment analysis and annotation. The node size represented the relative log_2_FC of each protein, while the edge width represented the interaction confidence.

### Weighted gene correlation network analysis (WGCNA) for proteomic data

When more than one Uniprot IDs match the same gene symbol (MGI symbol), the one with the lower unique peptide count or normalized abundance was removed. Then, two independent proteomic datasets were integrated with batch-effect correction by the HarmonizR R package with the default ComBat method ^111^. Confirmed by principal component analysis, group distribution within the same batch was preserved while group difference between batches was reduced. One NMNAT2 cKO; SARM1 WT sample was excluded as an outlier, and the remaining samples’ commonly detected proteins were analyzed by the R package WGCNA ^112^. The unsigned Pearson correlation matrix was calculated to measure gene co-expression similarity, and a weighted adjacency matrix was calculated by raising the correlation matrix to the power of a soft threshold equals 9. The minimum height (dissimilarity) for merging modules was set at 0.25. To define sample traits, samples with neurodegeneration were given 1, otherwise 0; samples with NMNAT2 expression were given 1, otherwise 0; SARM1 WT homozygous samples were given 2, heterozygous as 1, and KO as 0. For module-trait correlation analysis, all samples were used for examining correlation to neurodegeneration; SARM1 WT & heterozygous samples were selected for examining correlation to NMNAT2 expression; NMNAT2 cKO samples were selected for examining correlation to SARM1 gene dosage. The Pearson correlation coefficient and correlation *p*-value of the first principal component of the module protein expression profile (eigenprotein) and trait values were calculated to identify significant modules (P<0.05) correlated to a particular trait. The genes within the module of interest were input to the STRING website for PPI visualization and pathway annotation.

### Metabolomic mass spectrometry analysis

Gas chromatography–mass spectrometry (GC–MS) and liquid chromatography–mass spectrometry (LC–MS) experiments, along with metabolomic analysis, were conducted at the Metabolomics Core Facility at the University of Utah (Salt Lake City, UT, USA) (https://uofuhealth.utah.edu/diabetes-metabolism-research-center/research/core-facilities).

### GC-MS Sample Extraction

Tissue metabolites were extracted by transferring each sample to 2.0 mL ceramic bead mill tubes (Qiagen Catalog Number 13116-50). To each sample, 450 μL of cold 90% methanol (MeOH) solution containing the internal standard d4-succinic acid (Sigma 293075) was added per 25 mg of tissue. Samples were homogenized using an OMNI Bead Ruptor 24 and then incubated at -20°C for 1 hour. Following incubation, samples were centrifuged at 20,000 x g for 10 minutes at 4°C. Subsequently, 600 μL of supernatant was transferred from each bead mill tube into labeled, fresh microcentrifuge tubes. Another internal standard, d27-myristic acid, was added to each sample. Pooled quality control samples were generated by extracting a portion of supernatant from each sample. Process blanks consisting solely of extraction solvent underwent the same procedural steps as the actual samples. Finally, samples were dried en vacuo.

### GC-MS

GC-MS analyses were conducted using an Agilent 5977b GC-MS MSD-HES coupled with an Agilent 7693A automatic liquid sampler. Dried samples were reconstituted in 40 µL of a 40 mg/mL solution of O-methoxylamine hydrochloride (MOX) in dry pyridine and incubated for one hour at 37 °C. Subsequently, 25 µL of this solution was transferred to auto sampler vials. An automatic addition of 60 µL of N-methyl-N-trimethylsilyltrifluoracetamide (MSTFA with 1% TMCS) was performed using the auto sampler, followed by incubation for 30 minutes at 37 °C. After incubation, samples were vortexed, and 1 µL of the prepared sample was injected into the gas chromatograph inlet in split mode with the inlet temperature maintained at 250°C. A split ratio of 5:1 was used for most metabolites; however, metabolites saturating the instrument at this ratio were analyzed using a 50:1 split. The gas chromatograph was programmed with an initial temperature of 60°C for one minute, followed by a ramp of 10°C/min to 325°C, and held for 10 minutes. Separation was achieved using a 30-meter Agilent Zorbax DB-5MS with 10 m Duraguard capillary column, with helium as the carrier gas at a flow rate of 1 mL/min.

### LC-MS Sample Extraction

Samples were extracted in 450 μL of ice-cold 4:1 methanol:water solution, supplemented with 0.1 μg/mL carnitine-d9 internal standard, 1 mg/mL N-ethylmaleimide, and 0.1% ammonium hydroxide. A process blank was prepared using 50 μL of ddH2O and 450 μL of the extraction solution, following the same extraction procedure. Homogenization of samples was carried out in a bead mill for 30 seconds, followed by transfer to new Eppendorf tubes and incubation at - 20°C for 1 hour. Subsequently, samples were centrifuged at 20,000 x g for 10 minutes at 4°C, and the resulting supernatant was transferred to a fresh Eppendorf tube for further analysis.

### LC-MS

Prior to analysis, both samples and process blanks (PBs) were transferred to PTFE autosampler vials. A pooled quality control (QC) sample was created by combining 5 µL from each individual sample. Before analysis, samples were randomized. Analysis was conducted using a SCIEX 6500 QTRAP coupled with a SCIEX Nexera UPLC system (AB SCIEX LLC, Framingham, MA, USA) operating in positive-mode multiple reaction monitoring (MRM). Separation was achieved using a Sequant ZIC-pHILIC 2.1 × 100 mm column (Millipore Sigma, Burlington, MA, USA) with a Phenomenex Krudkatcher (Phenomenex, Torrance, CA, USA). The chromatographic gradient started with an initial concentration of 99% ACN with 5% ddH2O (Buffer B) and 5% 25 mM ammonium carbonate in ddH2O (Buffer A), held for 3.3 minutes at a flow rate of 0.15 mL/min. Buffer B was then decreased to 15% over 7.5 minutes and held for 3.8 minutes. Finally, Buffer B was returned to its initial concentration over 0.1 minute, and the system was allowed to re-equilibrate for 15 minutes between runs.

### MS data Analysis

The GC-MS data were collected using MassHunter software (Agilent), while metabolites were identified and their peak areas recorded using MassHunter Quant. LC-MS data were collected and analyzed using SCIEX MultiQuant software. Metabolite identities were determined using transitions from Metlin (https://metlin.scripps.edu/landing_page.php?pgcontent=mainPage) and retention time data from in-house pure standards. Subsequently, the data were transferred to a Microsoft Excel spreadsheet. Metabolite identity was confirmed using a combination of an in-house metabolite library developed with pure purchased standards, the NIST library, and the Fiehn library. Statistical analysis and pathway mapping were performed using MetaboAnalyst (https://www.metaboanalyst.ca/) ^113^.

### Proteomic mass spectrometry analysis

The proteomic mass spectrometry analysis conducted in this study was carried out by the Indiana University School of Medicine Proteomics Core (Indianapolis, IN, USA) (https://medicine.iu.edu/service-cores/facilities/proteomics).

### Proteomic sample preparation

The flash-frozen tissue samples were crushed using a CryoPrep homogenizer (Covaris™). Subsequently, the samples were centrifuged at 14,000 x g for 20 minutes, and protein concentrations were determined using the Bradford protein assay (BioRad Cat No: 5000006). Each sample containing 50 µg equivalent of protein underwent treatment with 5 mM tris(2-carboxyethyl) phosphine hydrochloride (Sigma-Aldrich Cat No: C4706) to reduce disulfide bonds, followed by alkylation of resulting free cysteine thiols with 10 mM chloroacetamide (Sigma Aldrich Cat No: C0267). The samples were then diluted with 50 mM Tris.HCl pH 8.5 (Sigma-Aldrich Cat No: 10812846001) to a final urea concentration of 2 M for overnight Trypsin/Lys-C digestion at 35 °C, using a 1:50 protease to substrate ratio (Mass Spectrometry grade, Promega Corporation, Cat No: V5072)

### Peptide Purification and Labeling

The digested samples were acidified with trifluoroacetic acid (TFA, 0.5% v/v) and subjected to desalting using Waters Sep-Pak® Vac cartridges (Waters™ Cat No: WAT054955), with a 1 mL wash of 0.1% TFA followed by elution in 0.6 mL of 70% acetonitrile with 0.1% formic acid (FA). The resulting peptides were dried using a speed vacuum and resuspended in 50 mM triethylammonium bicarbonate. Subsequently, peptide concentrations were assessed using the Pierce Quantitative colorimetric assay (Cat No: 23275). Equal amounts of peptide from each sample were labeled with 0.5 mg of Tandem Mass Tag Pro (TMTpro) reagent (16-plex kit, Thermo Fisher Scientific, TMTpro™ Isobaric Label Reagent Set; Cat No: 44520, lot no. YG370070 and YI372118, according to the manufacturer’s instructions (Li et al., 2020). After confirming >98% labeling efficiency, reactions were quenched with 0.3% hydroxylamine (v/v) at room temperature for 15 minutes. The labeled peptides were then combined, dried using a speed vacuum, and desalted to remove excess label using a 100 mg Waters SepPak cartridge, eluted in 70% acetonitrile with 0.1% formic acid, and lyophilized to dryness.

### High pH Basic Fractionation

The combined samples were resuspended in 10 mM ammonium formate at pH 10. Half of each mixture underwent fractionation using an ofline Thermo UltiMate 3000 HPLC system equipped with a Waters Xbridge C18 column (3.5 µm x 4.6 mm x 250 mm, Cat No: 186003943). The elution gradient consisted of Buffer A (10 mM formate at pH 10) and Buffer B (95% acetonitrile with 10 mM formate at pH 10) at a flow rate of 1 mL/min: 0-15% B over 5 min, 15-20% B over 5 min, 20-35% B over 75 min, 35-50% B over 5 min, and 50-60% B over 10 min, followed by a 6-minute hold at 60% B. Fractions were collected continuously every 60 seconds into 96-well plates. Initial and late fractions with minimal material were combined and lyophilized, while the remaining fractions were concatenated into 24 fractions, dried, and resuspended in 50 µL of 0.1% formic acid prior to online LC-MS analysis.

### Nano-LC-MS/MS

Mass spectrometry analysis was conducted using an EASY-nLC 1200 HPLC system (SCR: 014993, Thermo Fisher Scientific) coupled with an Exploris 480™ mass spectrometer featuring a FAIMSpro interface (Thermo Fisher Scientific). One-fifth of each fraction was injected onto a 25 cm Aurora column (Ion Opticks) at a flow rate of 350 nL/min. The chromatographic gradient was composed of mobile phases A (0.1% formic acid (FA) in water) and B (0.1% FA, 80% acetonitrile (Thermo Fisher Scientific Cat No: LS122500)). The gradient increased from 8% to 38% B over 98 minutes, followed by a 10-minute ramp to 80% B, held for 2 minutes, and then decreased from 80% to 4% B over the final 5 minutes. Mass spectrometry was conducted in positive ion mode with a default charge state of 2, employing advanced peak determination and a lock mass of 445.12003. Three FAIMS CVs (-45, -55, and -65 CV) were utilized, each with a cycle time of 1 second and identical MS and MS2 parameters. Precursor scans (m/z 375-1500) were performed at an orbitrap resolution of 120,000, with an RF lens% of 40, automatic maximum injection time, standard AGC target, and a minimum MS2 intensity threshold of 5e3. Dynamic exclusion was set at 30 seconds. MS2 scans were conducted with a quadrupole isolation window of 1.6 m/z, 30% HCD collision energy, a resolution of 15,000, standard AGC target, automatic maximum injection time, and a fixed first mass of 110 m/z.

### Proteomic mass spectrometry Data Analysis

Resulting RAW files were analyzed in Proteome Discover™ 2.5 (Thermo Fisher Scientific) with a Mus musculus UniProt reference proteome FASTA (downloaded 022823, 55250 sequences) plus common laboratory contaminants (73). SEQUEST HT searches were conducted with full trypsin digest, 3 maximum number missed cleavages; precursor mass tolerance of 10 ppm; and a fragment mass tolerance of 0.02 Da. Static modifications used for the search were, 1) carbamidomethylation on cysteine (C) residues; 2) TMTpro label on lysine (K) residues. Dynamic modifications used for the search were TMTpro label on N-termini of peptides, oxidation of methionines, deamidation of asparagine or arginine, phosphorylation on serine, threonine or tyrosine, and acetylation, methionine loss or acetylation with methionine loss on protein N-termini. Percolator False Discovery Rate was set to a strict setting of 0.01 and a relaxed setting of 0.05. IMP-ptm-RS node was used for all modification site localization scores. Values from both unique and razor peptides were used for quantification. In the consensus workflows, peptides were normalized by total peptide amount with no scaling. Quantification methods utilized TMTpro isotopic impurity levels available from Thermo Fisher Scientific. Reporter ion quantification was allowed with S/N threshold of 6 and co-isolation threshold of 30%. Resulting grouped abundance values for each sample type, abundance ratio values; and respective p-values (t-test) from Proteome Discover were exported to Microsoft Excel.

### Lipidomic mass spectrometry analysis

The lipidomic mass spectrometry analysis conducted in this study was carried out by Metware Biotechnology Inc. (Woburn, MA, USA) (www.metwarebio.com). Ultra-performance liquid chromatography-tandem mass spectrometry (UPLC-MS/MS) was performed for qualitative and quantitative compound analysis.

### Lipidomic sample preparation

Take out the sample from the -80℃ refrigerator and thaw it on ice. Multi-point sample and weigh 20 mg of sample, homogenize (30 HZ) for 20 s with a steel ball and the centrifuge (3000 rpm, 4°C) for 30 s. Then add 1mL of the extraction solvent (MTBE: MeOH =3:1, v/v) containing internal standard mixture. After whirling the mixture for 15 min, 200 μL of ultrapure water was added. Vortex for 1 min and centrifuge at 12,000 rpm for 10 min. 200 μL of the upper organic layer was collected and evaporated using a vacuum concentrator. The dry extract was dissolved in 200 μL reconstituted solution (ACN: IPA=1:1, v/v) to LC-MS/MS analysis.

### Chromatography-mass spectrometry acquisition conditions

The data acquisition instruments consisted of Ultra Performance Liquid Chromatography (UPLC) (Nexera LC-40, https://www.shimadzu.com) and tandem mass spectrometry (MS/MS) (Triple Quad 6500+, https://sciex.com/ ).

#### Liquid phase conditions

1. Chromatographic column: Thermo Accucore™C30 (2.6 μm, 2.1 mm×100 mm i.d.)
2. Mobile phase: A phase was acetonitrile /water (60/40, V/V) (0.1% formic acid added, 10 mmol/L ammonium formate); B phase was acetonitrile / Isopropyl alcohol (10/90, V/V) (0.1% formic acid added, 10 mmol/L ammonium formate)
3. Gradient program: 80:20(V/V) at 0 min, 70:30(V/V) at 2 min, 40:60(V/V) at 4 min , 15:85(V/V) at 9 min, 10:90(V/V) at 14 min, 5:95(V/V) at 15.5 min, 5:95(V/V) at 17.3 min, 80:20(V/V) at 17.5 min, 80:20(V/V) at 20 min;
4. Flow rate: 0.35 ml/min; Column temperature: 45℃; Injection volume: 2 μL.

#### Mass spectrometry conditions

LIT and triple quadrupole (QQQ) scans were acquired on a triple quadrupole-linear ion trap mass spectrometer (QTRAP), QTRAP® 6500+ LC-MS/MS System, equipped with an ESI Turbo Ion-Spray interface, operating in positive and negative ion mode and controlled by Analyst 1.6.3 software (Sciex). The ESI source operation parameters were as follows: ion source, turbo spray; source temperature 500 ℃; ion spray voltage (IS) 5500 V(Positive),-4500 V (Neagtive); Ion source gas 1 (GS1), gas 2 (GS2), curtain gas (CUR) were set at 45, 55, and 35 psi, respectively.

Instrument tuning and mass calibration were performed with 10 and 100 μmol/L polypropylene glycol solutions in QQQ and LIT modes, respectively. QQQ scans were acquired as MRM experiments with collision gas (nitrogen) set to 5 psi. DP and CE for individual MRM transitions was done with further DP and CE optimization. A specific set of MRM transitions were monitored for each period according to the lipids eluted within this period.

### Principles of lipid qualification and quantification

With the in-house database MWDB (www.metwarebio.com), lipids were annotated based on its retention time and ion-pair information from MRM mode. In MRM mode, the first quadrupole screens the precursor ions for the target substance and excludes ions of other molecular weights. After ionization induced by the impact chamber, the precursor ions were fragmented, and a characteristic fragment ion was selected through the third quadrupole to exclude the interference of other non-target ions. By selecting a particular fragment, quantification is more accurate and reproducible.

