## Supplementary data for "NAD^+^ reduction in glutamatergic neurons triggers fatty acid catabolism and neuroinflammation in the brain, mitigated by SARM1 deletion"

**Sup. Figure 1**

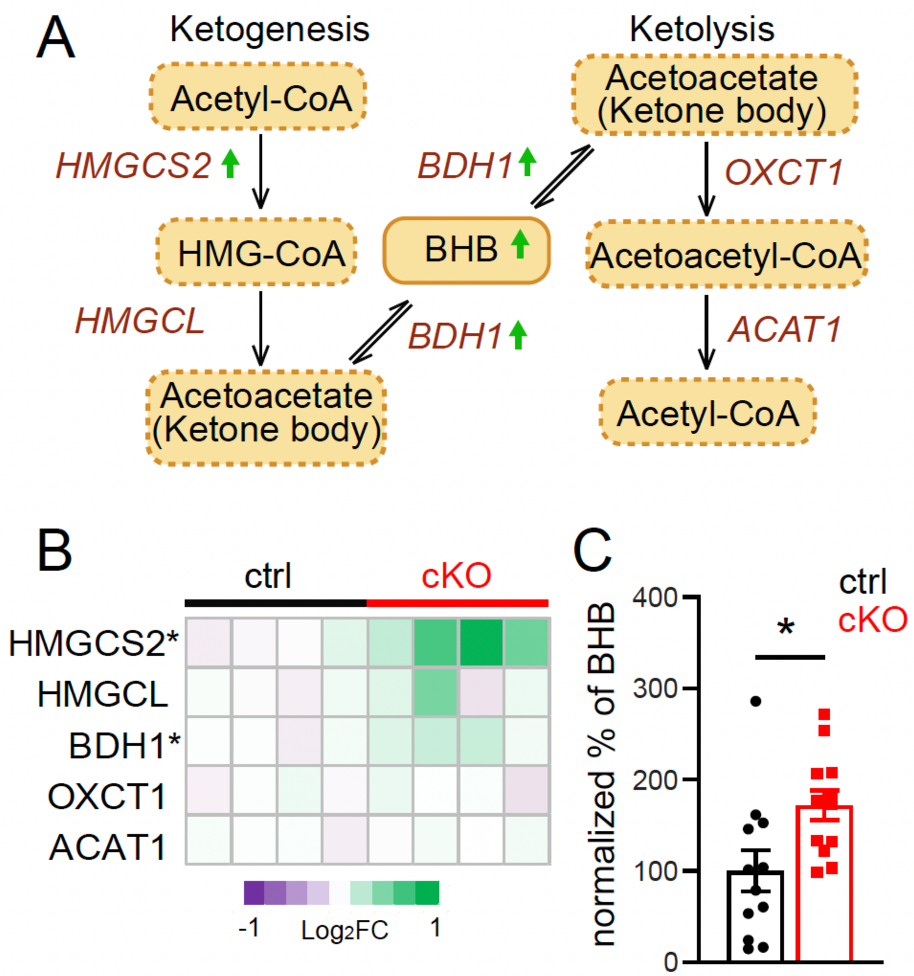


**Sup Fig. 1** Altered ketogenesis and ketolysis in NMNAT2 cKO cortices.

**(A)** Scheme of ketogenesis and ketolysis pathway. **(B)** Heatmaps of log2-transformed values summarize the enzymes of ketone metabolism (n=4 males per group). **(C)** Summary of normalized percentage of 3-Hydroxybutyric acid (BHB). Sample size: P16-P18 ctrl and cKO, n = 6 per sex. *, $p<0.05$ by Student's t-test. Metabolites labeled with a dashed line were not detected in the MS analysis. Abbreviations: ACAT1, acetyl-CoA acetyltransferase; BDH1, D-beta-hydroxybutyrate dehydrogenase; HMGCL, hydroxymethylglutaryl-CoA lyase; HMGCS2, hydroxymethylglutaryl-CoA synthase; OXCT1, succinyl-CoA:3-ketoacid coenzyme A transferase 1.

**Sup. Figure 2**

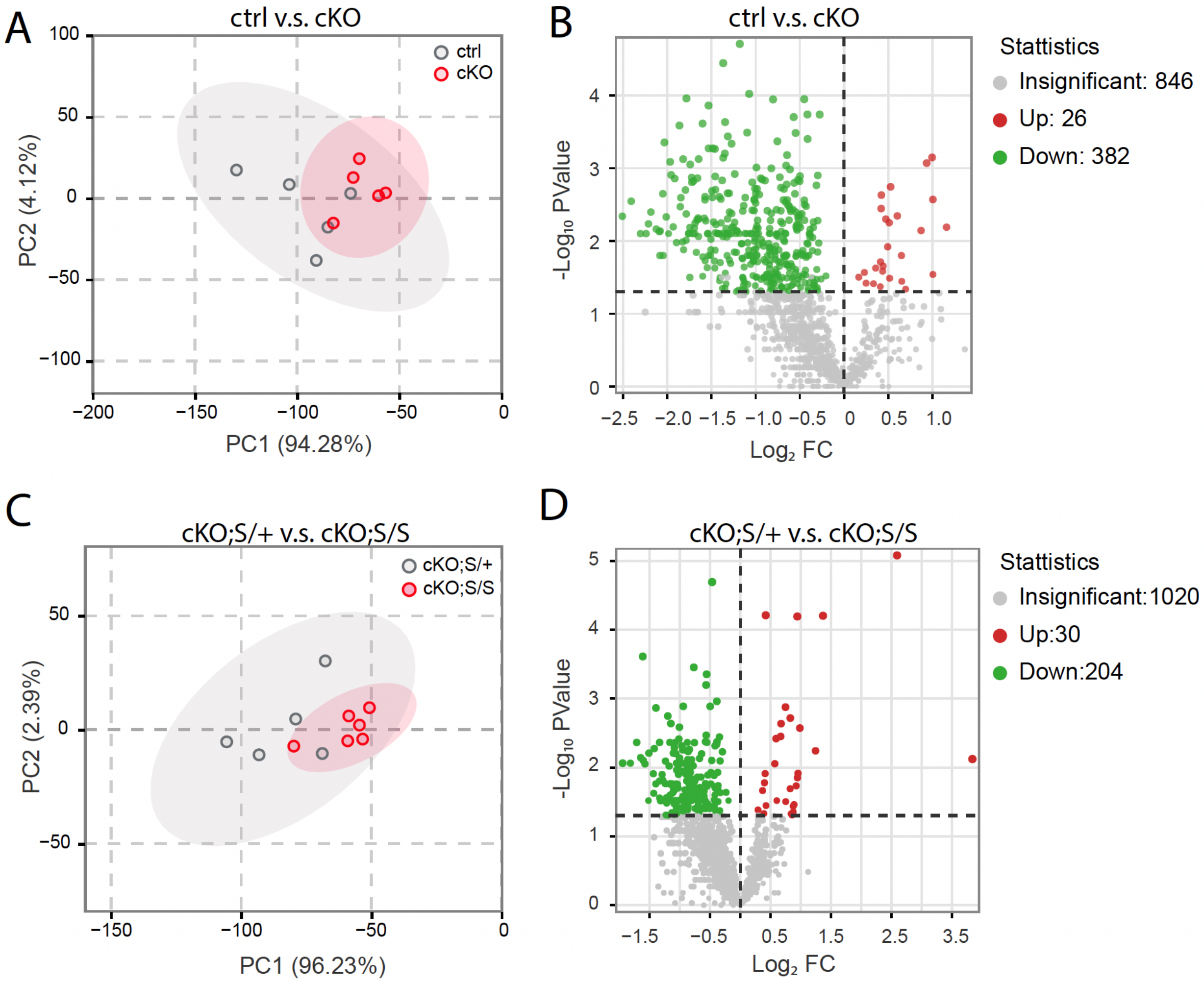


**Sup Fig. 2** Summary of lipidomic analyses. **(A,C)** Principal component analysis (PCA) reveals distinct separation of lipidomic profiles among genotypes. **(B,D)** Volcano plots show differentially expressed lipids upon NMNAT2 loss and the extent of recovery following SARM1 deletion.

**Sup. Figure 3**

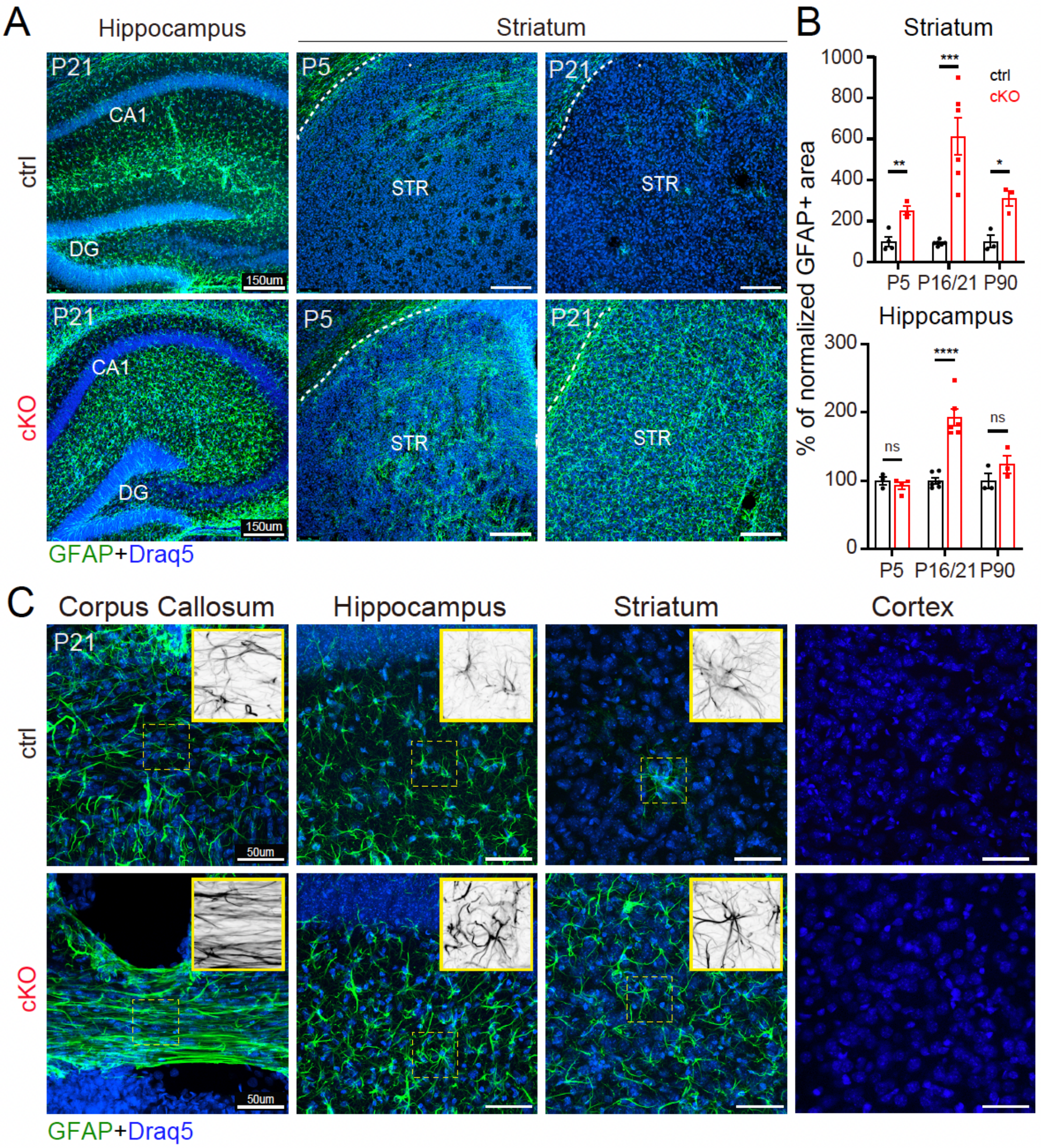


**Sup Fig. 3** Neuronal NMNAT2 loss increases astrocyte reactivities upon. **(A)** Representative GFAP staining images from P21 ctrl and NMNAT2 cKO hippocampus and striatum. **(B)** Bar graphs summarize normalized % of GFAP+ area at P4/5 (ctrl, n=3; cKO, n=4), P16/21 (n=6 per group), and P90 (n=3 per group) **(C)** High-magnification images with enlarged views (inserts) showing morphology of GFAP positive astrocytes in P21 corpus callosum, hippocampus, striatum, and cortex of ctrl and cKO brains. Almost no GFAP signals are present in upper cortical regions. Draq5 signals reveal nuclei locations. *, p<0.05; **, p<0.01; ***, p<0.001; ****, p<0.0001 by Student's t-test.

**Sup. Figure 4**

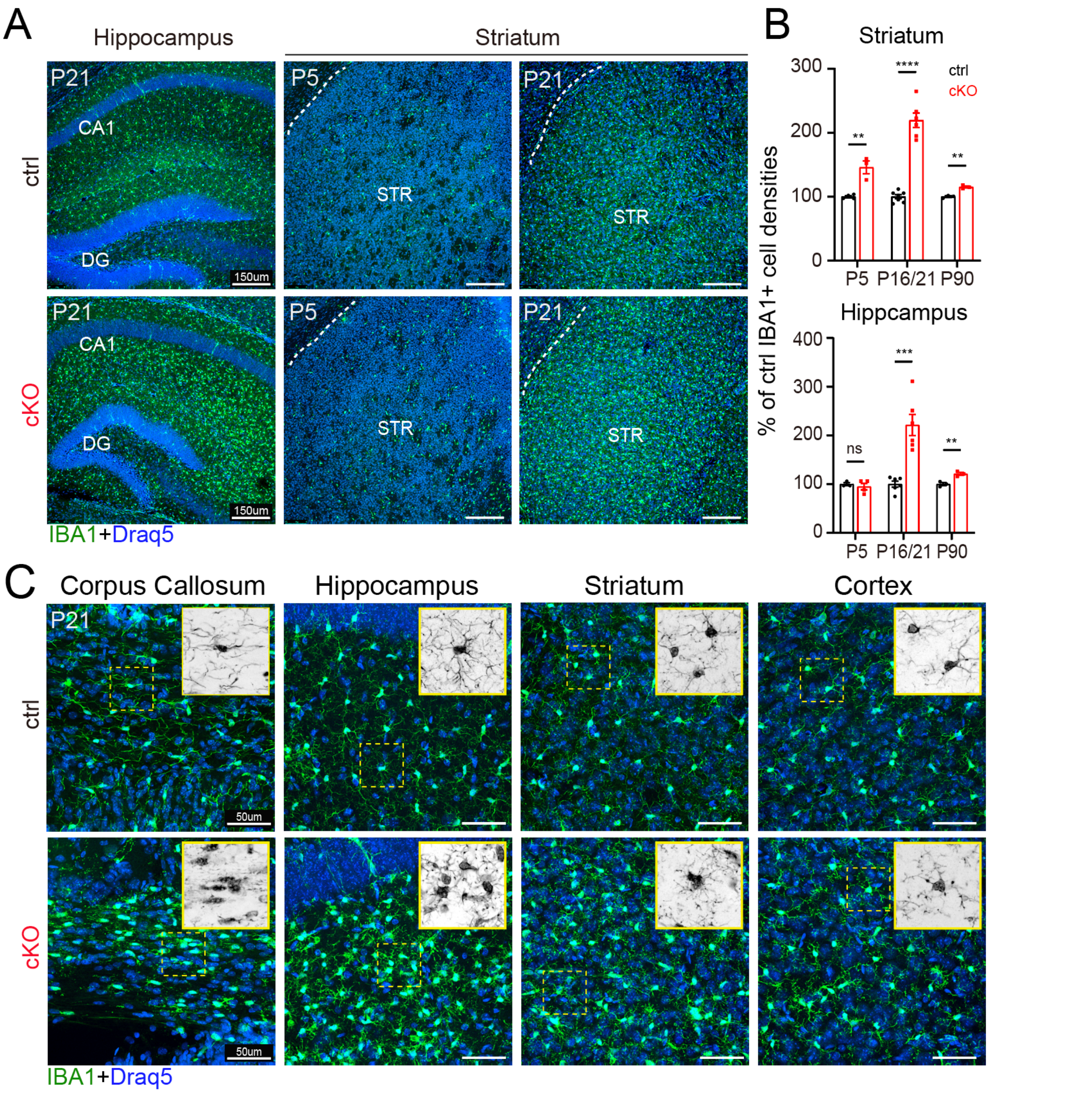


**Sup Fig. 4** Neuronal NMNAT2 loss increases the number of microglia and reactivities. **(A)** Representative IBA1 staining images from the hippocampus and striatum of NMNAT2 cKO and littermate control mice. **(B)** Bar graphs summarize the percentage of IBA1+ densities to controls at P4/5 (ctrl, n=3; cKO, n=4), P16/21 (n=6 per group), and P90 (n=3 per group). **(C)** High magnification images with enlarged views (inserts) showing IBA1+ microglia morphology in P21 corpus callosum, hippocampus, striatum, and cortex of control and cKO brains. *, p<0.05; **, p<0.01; ***, p<0.001; ****, p<0.0001 by Student's t-test.

**Sup. Figure 5**

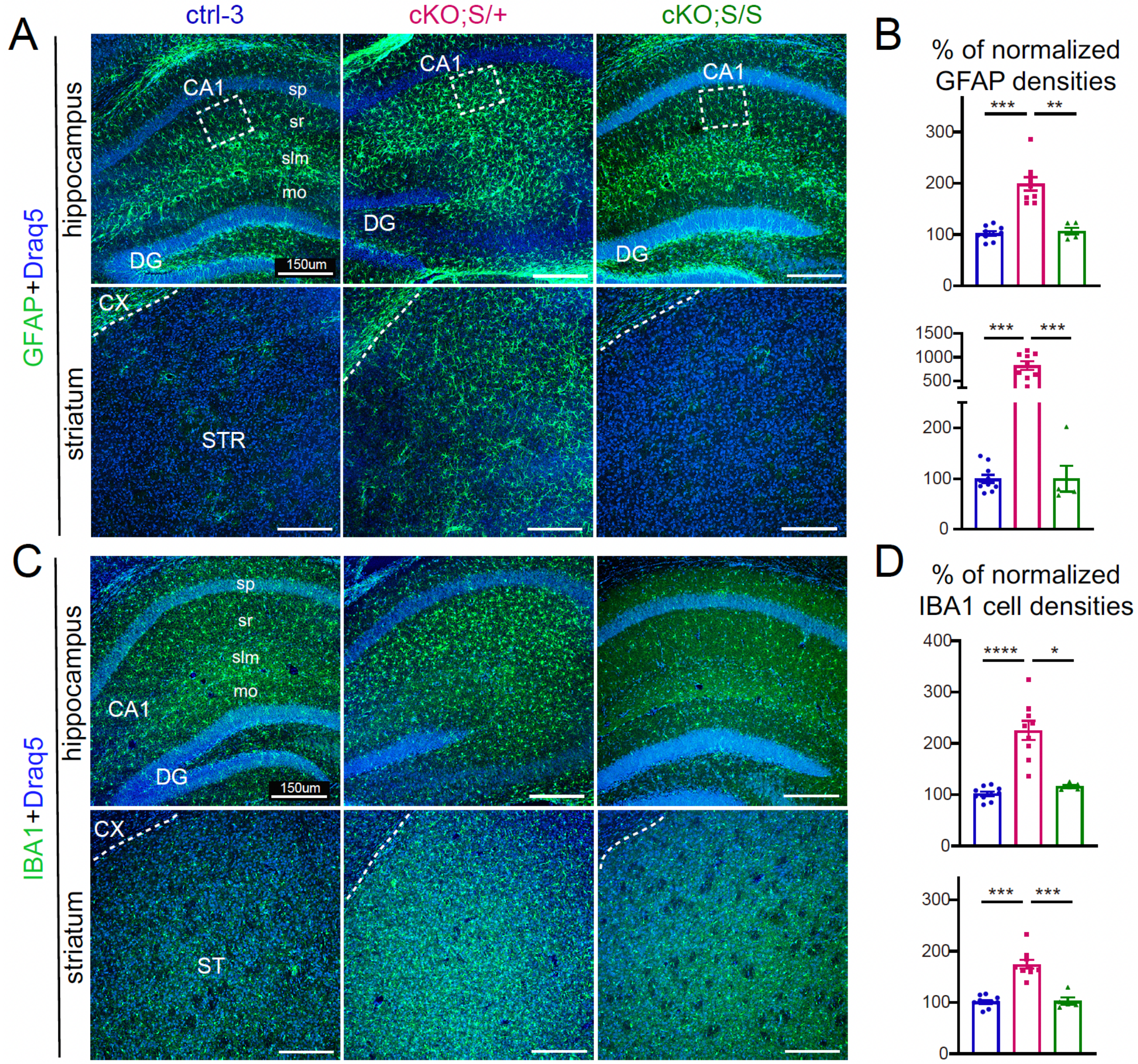


**Sup Fig. 5** Complete SARM1 deletion ameliorates neuroinflammatory responses in NMNAT2 cKO brains. (**A**) Representative GFAP staining images in the hippocampus and striatum of the three genotypes. (**B**) Summary of normalized % of GFAP+ densities in hippocampus and striatum (control-3, n=10; cKO;$S^{null/+}$, n=9; cKO;$S^{null/null}$, n=5). (**C**) Representative IBA1 staining images from the hippocampus and striatum at P21. (**D**) Summary of normalized % of IBA1+ cell densities in hippocampus and striatum (ctrl, n=10; cKO;$S^{null/+}$, n=9; cKO;$S^{null/null}$, n=5) *, p<0.05; **, p<0.01; ***, p<0.001; ****, p<0.0001 by Kruskal-Wallis test. Abbreviations: ctrl-3, control-3; cKO;S/+, cKO;$S^{null/+}$; cKO;S/+, cKO;$S^{null/null}$.

### Supplementary tables

**Sup. Table 1**. The list of fold changes of individual lipid subclasses for cKO to control and cKO;S$^{null/+}$ to cKO;S$^{null/null}$ cortices. Arrows indicate the direction of regulation (upregulation or downregulation). *, p<0.05; **, p<0.01; ***, p<0.001; ****, p<0.0001 by Student's t-test. 4 male mice per genotype per group.

| Lipid class | | cKO vs ctrl | cKO;Snull/+ vs. cKO;Snull/Snull |
|---|---|---|---|
| BA | Bile Acids | ▲ 1.78** | ▲ 1.28* |
| BMP | Bis(monoacylglycerol)phosphate | ns | ns |
| CAR | Acylcarnitine | ns | ns |
| Cer-ADS | Ceramide alpha-hydroxy fatty acid dihydrosphingosine | ▼ 0.37* | ns |
| Cer-AP | Ceramide alpha-hydroxy fatty acid phytosphingosine | ▼ 0.36** | ▼ 0.49* |
| Cer-AS | Ceramide alpha-hydroxy fatty acid sphingosine | ▼ 0.61* | ns |
| Cer-NDS | Ceramide non-hydroxy fatty acid dihydrosphingosine | ns | ns |
| Cer-NP | Ceramide non-hydroxy fatty acid phytosphingosine | ▼ 0.31** | ▼ 0.45* |
| Cer-NS | Ceramide non-hydroxy fatty acid sphingosine | ns | ▼ 0.66* |
| CerP | Ceramide-1-phosphate | ns | ▲ 1.82* |
| Cholesterol | Cholesterol | ns | ns |
| CoQ | Coenzyme Q | ns | ns |
| DG | Diacylglycerol | ns | ns |
| DG-O | Alkyl-diacylglycerol | ns | ns |
| DGDG | Digalactosyldiacylglycerol | ns | ns |
| Eicosanoid | Eicosanoids | ns | ns |
| FFA | Free Fatty Acid | ▼ 0.70*** | ▼ 0.79** |
| Hex2Cer | Dihexosylceramide | ns | ns |
| HexCer-AP | Hexosylceramide alpha-hydroxy fatty acid phytosphingosine | ▼ 0.58* | ns |
| HexCer-NS | Hexosylceramide non-hydroxy fatty acid sphingosine | ▼ 0.49* | ▼ 0.58* |
| LNAPE | Lyso-N-acyl phosphatidylethanolamine | ns | ns |
| LPC | Lysophosphatidylcholine | ▼ 0.71*** | ns |
| LPC-O | Alkyl-lysophosphatidylcholine | ns | ns |
| LPE | Lysophosphatidylethanolamine | ▼ 0.76** | ns |
| LPE-P | Alkenyl-lysophosphatidylethanolamine | ns | ns |
| LPG | Lysophosphatidylglycerol | ▼ 0.77* | ns |
| LPI | Lysophosphatidylinositol | ▼ 0.58** | ▼ 0.70*** |
| LPS | Lysophosphatidylserine | ▼ 0.67* | ns |
| MG | Monoacylglycerol | ns | ▲ 2.25**** |
| MGDG | Monogalactosyldiacylglycerol | ▼ 0.52* | ns |
| PA | Phosphatidic Acid | ns | ns |
| PC | Phosphatidylcholine | ns | ns |
| PC-O | Ether-linked Phosphatidylcholine | ns | ns |
| PE | Phosphatidylethanolamine | ns | ns |
| PE-O | Ether-linked Phosphatidylethanolamine | ▼ 0.82* | ns |
| PE-P | Plasmalogen Phosphatidylethanolamine | ns | ns |
| PG | Phosphatidylglycerol | ns | ns |
| PI | Phosphatidylinositol | ▼ 0.66* | ns |
| PMeOH | Phosphatidylmethanol | ns | ns |
| PS | Phosphatidylserine | ▼ 0.62* | ns |
| SHexCer | Sulfatide (Sulfated Hexosylceramide) | ns | ns |
| SM | Sphingomyelin | ▼ 0.75* | ns |
| SPH | Sphingosine | ns | ns |
| TG | Triacylglycerol | ns | ns |

**Sup. Table 2**. The top 15 KEGG pathways enrichment analysis with significantly altered lipids. Pathways are ranked by the number of significant lipid compounds detected. 4 male mice per genotype per group.

| cKO vs ctrl | | | |
|---|---|---|---|
| **Kegg_pathway** | **ko_ID** | **total #** | **%** |
| Glycerophospholipid metabolism | ko00564 | 511 | 35.81 |
| Sphingolipid metabolism | ko00600 | 268 | 36.19 |
| Ether lipid metabolism | ko00565 | 219 | 39.27 |
| Glycosylphosphatidylinositol (GPI)-anchor biosynthesis | ko00563 | 154 | 38.96 |
| Autophagy - animal | ko04140 | 154 | 38.96 |
| Retrograde endocannabinoid signaling | ko04723 | 207 | 28.99 |
| Choline metabolism in cancer | ko05231 | 201 | 28.36 |
| Sphingolipid signaling pathway | ko04071 | 145 | 33.79 |
| Leishmaniasis | ko05140 | 121 | 38.84 |
| Necroptosis | ko04217 | 116 | 35.34 |
| Kaposi sarcoma-associated herpesvirus infection | ko05167 | 99 | 39.39 |
| Insulin resistance | ko04931 | 223 | 17.49 |
| Adipocytokine signaling pathway | ko04920 | 96 | 30.21 |
| Diabetic cardiomyopathy | ko05415 | 97 | 29.90 |
| Neurotrophin signaling pathway | ko04722 | 85 | 30.59 |

| cKO;S/+ compared to cKO;S/S | | | |
|---|---|---|---|
| **Kegg_pathway** | **ko_ID** | **total #** | **%** |
| Sphingolipid metabolism | ko00600 | 268 | 34.70 |
| Glycerophospholipid metabolism | ko00564 | 511 | 16.05 |
| Sphingolipid signaling pathway | ko04071 | 145 | 33.79 |
| Insulin resistance | ko04931 | 223 | 21.52 |
| Leishmaniasis | ko05140 | 121 | 38.84 |
| Necroptosis | ko04217 | 116 | 36.21 |
| Adipocytokine signaling pathway | ko04920 | 96 | 43.75 |
| Diabetic cardiomyopathy | ko05415 | 97 | 43.30 |
| Neurotrophin signaling pathway | ko04722 | 85 | 45.88 |
| Ether lipid metabolism | ko00565 | 219 | 13.24 |
| Retrograde endocannabinoid signaling | ko04723 | 207 | 13.53 |
| Choline metabolism in cancer | ko05231 | 201 | 11.94 |
| Glycosylphosphatidylinositol (GPI)-anchor biosynthesis | ko00563 | 154 | 14.94 |
| Autophagy - animal | ko04140 | 154 | 14.94 |
| Kaposi sarcoma-associated herpesvirus infection | ko05167 | 99 | 20.20 |

**Sup. Table 3**. Summary for proteomic changes on selected proteins involved in lipid metabolism, inflammation responses, and glutathione metabolism. Arrows indicate the direction of regulation. *, p<0.05; **, p<0.01; ***, p<0.001 by Student's t-test.

| Lipid Metabolism | | | |
|---|---|---|---|
| | Gene ID | cKO vs ctrl | cKO;S/+ vs cKO;S/S |
| *Fatty Acid Synthesis and Modification* | HSD17B11 | ▲ 1.135* | ns |
| | HSD17B12 | ▼ 0.809** | ns |
| | ACSF2 | ▲ 1.164* | ns |
| | ACSL3 | ▲ 1.175*** | ns |
| | ECI2 | ▲ 1.115* | ns |
| | OXSM | ▲ 1.073** | ns |
| | AACS | ▼ 0.675*** | ▼ 0.797** |
| *Lipogenes is* | ACACA | ▼ 0.899* | ns |
| | FASN | ▼ 0.929* | ns |
| | ACAT1 | ns | ns |
| | ACSS2 | ▼ 0.776*** | ▼ 0.874** |
| *Fatty Acid β-Oxidation* | ACADVL | ▲ 1.145* | ns |
| | CPT2 | ▲ 1.304* | ns |
| | HADHB | ▲ 1.159* | ns |
| | DECR1 | ▲ 1.133* | ns |
| | SCCPDH | ▼ 0.875* | ns |
| *Ketogene sis* | HMGCS2 | ▲ 1.588* | ns |
| | HMGCL | ns | ns |
| | BDH1 | ▲ 1.118* | ns |
| | OXCT1 | ns | ns |
| *Cholesterol and Steroid Metabolism* | ABCA1 | ▲ 1.966** | ns |
| | APOE | ▲ 1.215** | ns |
| | DHCR24 | ▼ 0.564** | ns |
| | HMGCR | ▼ 0.542** | ns |
| | HMGCS1 | ▼ 0.575** | ▼ 0.599** |
| | FDFT1 | ▼ 0.678** | ns |
| | FDPS | ▼ 0.843* | ns |
| | PMVK | ▼ 0.706** | ▼ 0.753** |
| *Sphingolipid Metabolism* | CERS2 | ▼ 0.771* | ns |
| | CERS4 | ▼ 0.584* | ns |
| | SMPD1 | ▼ 0.812** | ns |
| | ST8SIA4 | ▼ 0.892* | ns |
| | HEXB | ▲ 1.194* | ns |
| | CPTP | ns | ▼ 0.927* |
| | CERT1 | ns | ▲ 1.090* |
| *Glycerophospholipid Metabolism* | LPCAT2 | ▼ 0.876* | ns |
| | LPGAT1 | ▼ 0.852** | ns |
| | PCYT2 | ▼ 0.874** | ns |
| | PTDSS2 | ▼ 0.798* | ns |
| | PAFAH1B3 | ▲ 1.402* | ns |
| | GDE1 | ns | ▲ 1.459* |
| | GPCPD1 | ns | ▲ 1.124* |
| | LCLAT1 | ns | ns |
| | LPCAT4 | ns | ns |
| | LPCAT1 | ns | ns |
| *Lipolysis and Lipid Signaling* | PLCD4 | ▲ 1.265* | ns |
| | PLD1 | ▼ 0.894* | ns |
| | PLD2 | ▲ 1.212* | ns |
| | PRKAG2 | ▼ 0.730* | ns |
| | PNPLA2 | ▲ 2.064** | NA |

| Inflammatory pathways | | | |
|---|---|---|---|
| | Gene ID | cKO vs ctrl | cKO;S/+ vs cKO;S/S |
| *Interleukin* | IL18 | ▲ 1.229* | ns |
| | IL1RAPL1 | ▲ 1.147* | ns |
| | IL1RAP | ns | NA |
| *Astrocytes and glial cell markers* | GFAP | ▲2.207*** | ▲2.057*** |
| | ALDH1L1 | ▲ 1.132* | ns |
| | S100B | ns | ns |
| | AIF1 | ns | ns |
| | AIF1L | ▼ 0.638* | ▼ 0.814** |
| | CD68 | ▲1.858*** | NA |
| Complement System | C1QTNF4 | ▲ 1.080** | ▲ 1.145** |
| | CD93 | ▼ 0.772* | ▼ 0.735** |
| | C1QB | ▲ 2.207** | ns |
| | C1QC | ▲ 2.649* | ns |
| | XRCC5 | ▲ 1.138* | ns |
| | C4B | ▲ 1.756** | ▲1.730*** |
| *Regulation of immune system process* | SDC4 | NA | ▲ 1.664** |
| | THBS4 | ▲ 1.525** | ▲ 2.313** |
| | CD47 | ns | ▲ 1.151* |
| | SIRPA | ns | ▲ 1.270* |
| | PTPN6 | ns | ▲ 1.160** |
| | MAPK1 | ns | ▲ 1.399* |

| Glutathione Metabolism | | | |
|---|---|---|---|
| | Gene ID | cKO vs ctrl | cKO;S/+ vs cKO;S/S |
| *Glutathione cycle* | GPX1 | ns | ns |
| | GPX7 | ns | ns |
| | GSR | ns | ns |
| *Glutathi one metabol* | GCLC | ns | ▲ 1.126* |
| | GCLM | ns | ns |
| | GSS | ns | ▲ 1.133* |
| Glutathione conjugation detoxification | GSTA4 | ns | ▲ 1.266* |
| | GSTK1 | ns | ns |
| | GSTM1 | ns | ▲ 1.304* |
| | GSTM2 | ns | ns |
| | GSTM4 | ▲ 1.198* | NA |
| | GSTM5 | ns | ns |
| | GSTM7 | ns | NA |
| *Mito chon drial* | SLC25A40 | ▼ 0.826* | NA |

**Sup. Table 4.** Lists of biological processes and proteins in Plum2 and Steelblue modules
Summary of MGCNA protein analysis and selected module characteristics. N= 4 male mice per genotype per group. cKOA refers to cKO;$S^{null/+}$ mice, and cKOB refers to cKO;$S^{null/Snull}$ mice.

**Sup. Table 5.** Statistics and numbers for figures and supplementary figures.
